# Aboveground enemy release increases seedling survival in grasslands

**DOI:** 10.1101/2023.10.06.561247

**Authors:** Joshua I. Brian, Harry E. R. Shepherd, María Ángeles Pérez-Navarro, Jane A. Catford

## Abstract

1. The enemy release hypothesis is a popular hypothesis to explain the success of invasive plants. Enemy release studies typically focus on single species or types of communities, feature indirect experimental manipulations that apply pesticides to whole communities not individual species, and only examine responses of established plants or plant populations, limiting their generality. Using a novel species-specific approach, we examine whether enemy release can enhance seedling survival and recruitment of 16 grassland species by experimentally linking enemy release with enhanced plant performance.
2. We planted seedlings of 16 native grassland species from two functional groups (C4 grasses and non-legume forbs) into two grassland sites (early and mid succession). We hand-painted 1,548 individual seedlings with pesticides (insecticide and fungicide) over the course of a growing season to enforce aboveground species-specific enemy release, and tested whether it enhanced survival relative to untreated controls. Using native species enabled us to directly test effects of enemy release, while avoiding confounding factors like unknown invasion histories. Of the 16 native study species, 13 are naturalised/invasive outside of their native ranges.
3. Release from insects increased seedling survival by 80% on average, with no additional benefit of release from fungal pathogens. This effect was consistent across functional groups and community successional stages, and was strongest in resource-acquisitive species. The size of species’ performance benefits from enemy release were positively correlated with the number of regions globally where each species has been introduced and naturalised.
4. *Synthesis*. Previous studies of enemy release have centred on adults and findings have varied among species. We found a positive effect of release from insect herbivores early in colonisation – a trend that held across functional groups and types of resident community. We posit that the consistent vulnerability of seedlings vis-à-vis later life stages leads to this more ubiquitous benefit of enemy release. Enemy release may therefore aid initial recruitment of most, if not all, plants during the invasion process, even if enemies rapidly accumulate. The positive correlations between the benefits of enemy release for seedlings, species’ life history strategies and global naturalisation patterns provide compelling hypotheses for future research.

## 1. Introduction

The enemy release hypothesis (ERH) is the best-known hypothesis explaining biological invasions (Enders *et al*., 2018). The ERH states that, on movement to a new range, species may escape the regulatory effects of enemies from their home range and can then become competitively dominant over native species in their invaded range (Keane & Crawley, 2002). The key mechanism of this dominance is that native species (in their home range) suffer enemy damage and thus remain regulated, while exotic species (in their invaded range) do not. However, evidence for when enemy release facilitates invasion is inconsistent and often contradictory (Jeschke & Heger, 2018; Brian & Catford, 2023). In particular, how enemies affect plant performance and the corresponding benefits of release remain understudied (Prior *et al*., 2015; Agrawal & Maron, 2022). Most studies focus on damage or diversity of enemies (Zhang *et al*., 2018b; Callaway *et al*., 2022), measures which may not directly translate to plant performance (Katz, 2016; Xu *et al*., 2021). Therefore, it is difficult to reliably link enemy release to the demographic advantages that are often exhibited by exotics (Brian & Catford, 2023). This absence of a causal link hinders interpretation of one of the key hypothesised causes of exotic species success.

A direct way to link enemy release with performance would be to enforce release by applying pesticides to individual plants and examining (potential) improvements in performance of released individuals. Enforcing enemy release does not ask whether it has occurred naturally, but whether it can explain increased performance when it does occur. We are aware of very few studies that use this method to directly link enemy release to plant performance in natural communities (but see Dewalt *et al*. 2004; Vasquez & Meyer 2011; Morris *et al*. 2022). Enforcing enemy release may be particularly powerful when native species are used as experimental subjects, as it avoids the confounding issue of invader minimum residence time and consequent variation in background levels of enemy release (Hawkes, 2007). Treating native individuals with pesticides in the field, and then comparing the performance of these individuals with non-treated individuals, tests the mechanism of enemy release: it releases specific individuals from enemy regulation, while co-occurring individuals in the community are still faced with enemies. Evidence for increased performance in released individuals relative to non-released individuals would then mechanistically support the link between enemy release and performance. This approach cannot determine whether enemy release has contributed to the success of a given invasive species, or the prevalence of enemy release among invasive species. However, it causally demonstrates that enemy release can be advantageous to individuals that are released, a link that has not been fully investigated in the enemy release literature (Brian & Catford 2023).

Experimental tests in field conditions are vital to contextualise the role of enemy release in invasion (Allen *et al*., 2020; Cuny *et al*., 2021), as plant performance is also affected by other factors such as environmental conditions and competition from native species. For example, even if not released from enemies, exotic plants that invade high resource environments could still experience increased performance if they compensate for any enemy-induced biomass losses through increased growth (Zhang *et al*., 2018b). An exotic that is phylogenetically closely related to native residents may faced increased competition due to high niche overlap, reducing any benefits from enemy release. However, studies on plant performance frequently take place in glasshouse environments (e.g. Dawson *et al*., 2014; Zhao *et al*., 2020), which differ from field conditions (Ojha *et al*., 2022). When enemy release has been experimentally tested in the field, treatments (e.g. pesticides) are typically applied over whole plots (e.g. Heckman *et al*., 2017; Suwa & Louda, 2012). Whole-plot applications assume that exotic plants naturally present in plots will not show a response to pesticide application (as they are already experiencing enemy release), while co-occuring native plants will benefit from pesticide application. However, plot-level applications cannot distentangle effects of enemy release from indirect effects of broad pesticide application such as changes in total plant richess or biomass (Agrawal & Maron, 2022), necessitating a targeted approach that releases specific species (Fig. S1).

Plant survival, particularly of seedlings, is an important performance metric determining the initial success and spread of exotic plants (Bohl Stricker & Stiling, 2013; Zhang *et al*., 2023). Juvenile plants are especially susceptible to enemies (Zhao *et al*., 2020; Bruns *et al*., 2022), so enemy release during plant recruitment may particularly affect outcomes of invasion. However, few empirical studies focus on enemy release during early recruitment (Hawkes, 2007) (but see Aldorfová *et al*. 2020), or on survival generally (Brian & Catford 2023), and many frameworks do not consider enemy release as important in the establishment phase of plant invasions (Colautti *et al*., 2004; Inderjit *et al*., 2005; Dawson *et al*., 2009). We therefore have scarce knowledge of whether and how enemy release increases exotic plant survival, especially in the initial colonisation phase (sensu Catford *et al*., 2009 and refs. therein).

The occurrence and strength of enemy release varies with the functional groups and traits of different exotic species. At a broad level, different functional groups experience pressure from different enemies. For example, the dominant enemy group of C4 grasses is fungal pathogens, while non-legume forbs are targeted more by insect herbivores (Ebeling *et al*., 2022). Release from enemies also appears to benefit non-legume forb biomass to a much greater extent than C4 grass biomass, a response that also depends on broader community diversity (Seabloom *et al*., 2018). Within functional groups, factors such as interspecific trait variation can also produce a wide variety of observed responses to enemy release. For example, species that invest heavily in growth at the expense of defence are more likely to suffer high levels of damage, and benefit the most from enemy release (Blumenthal *et al*., 2009). Despite this wide range of potential variation in plant responses to enemy release across functional groups and species, most studies in field conditions focus on one or two plant species (e.g. Louda & Potvin 1995; Dewalt *et al*. 2004; Eckberg *et al*. 2014), limiting the scope of potential inferences. To increase understanding of enemy release in early plant survival, there is a need to study a broad range of plant and enemy species from multiple functional groups.

In this study, we carried out a grassland field experiment to test the importance of enemy release for seedling survival over one growing season. We studied 16 native plant species from two functional groups (C4 grasses and non-legume forbs) released from two enemy guilds (aboveground fungal pathogens and aboveground insect herbivores), and examined the effects of release in communities at two different successional stages. 13 of our 16 target species have naturalised outside of their native range elsewhere in the world. We applied pesticides by hand-painting individual target plants, rather than spraying the whole community in a target plot, to enforce species-specific enemy release. This enabled us to directly test whether enemy release increases seedling survival and – if so – of which species and in which communities. We recognise that exotic species may be systematically different to natives (van Kleunen *et al*., 2010a), and so our experimental design targeting native species is not directly analagous to the arrival of exotic species. Nevertheless, by experimentally simulating the core mechanism of enemy release (an individual recruiting enemy-free into an enemy-regulated community), our experiment provides causal insight into how enemy release may increase performance and thus invasive potential.

## 2. Materials and Methods

### 2.1. Study site and species

Our experiment took place at Cedar Creek Ecosystem Science Reserve (hereon Cedar Creek) in Minnesota, USA. Cedar Creek has nitrogen-limited sandy soils, annual precipitation of ∼780 mm and mean annual temperature of 6.72 °C (averaged from 1987 to 2016 at Cedar Creek weather station). We studied 16 species, eight C4 grasses and eight non-legume forbs (Table 1). We chose these functional groups as they have previously displayed the most divergent responses to enemy removal (Seabloom *et al*., 2018). The eight species within each group were selected to cover a broad range of trait space (Fig. S2). All 16 species are native to Minnesota and present at Cedar Creek.

**Table 1:**
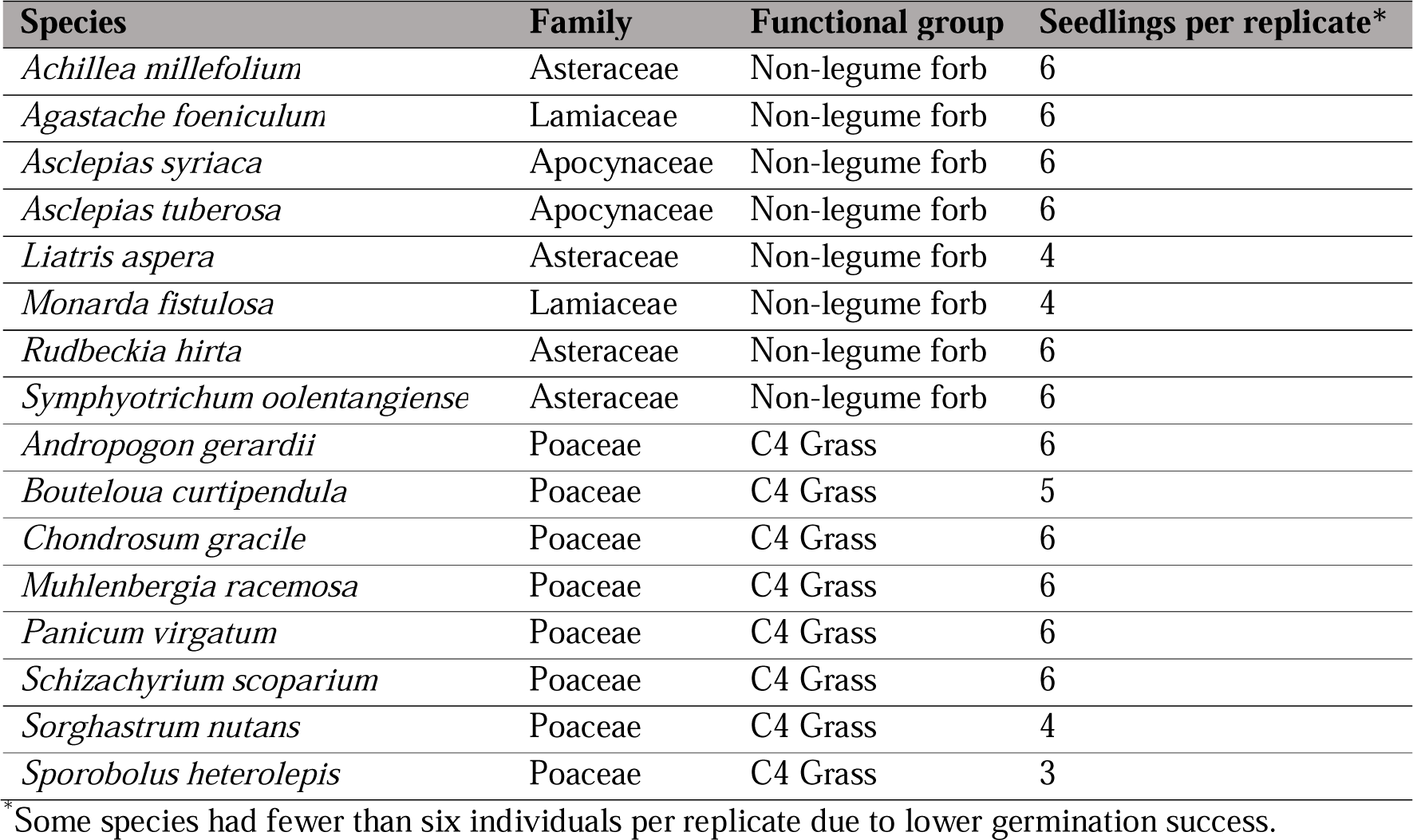
Study species in the experiment.

We sourced seeds from two native seed suppliers who collect seeds from local grassland populations (Prairie Moon, prairiemoon.com; Prairie Restorations, prairieresto.com). Individual seeds were planted in 12 cm deep ‘cone-tainers’, which were filled with sterilised soil (heated for 24 h at 80 °C) taken from Cedar Creek with a thin layer of vermiculite added on top. These were grown in a glasshouse for one month and irrigated regularly, after which time the seeds of all 16 species had germinated. The glasshouse was completely enclosed and monitored regularly for pests, protecting seedlings from enemies throughout germination and initial growth.

### 2.2. Experimental design

We established our experiment in the Lawrence Strips (45.414° N, -93.184° W), an old field at Cedar Creek approximately 100 m wide and 650 m long. This field was incrementally abandoned from agriculture: beginning in 1974, a 100 m × 20 m strip was abandoned each year, moving north through the field with the final strip abandoned in 2008 (Inouye *et al*., 1994). The field therefore contains a gradient of plant communities at different successional stages. We established plots in two strips in 2022: ‘early succession’ (20 years old, abandoned 2002) and ‘mid succession’ (41 years old, abandoned 1981) (Fig. S3). The diversity and composition of the two successional stages differed. Average summed plant cover was 55.8% in early plots and 75.4% in mid plots; early plots averaged 9.9 species per plot (range: 4-15) while mid plots averaged 12.2 species per plot (range: 6-18). The five most common species in early plots were *Digitaria cognata, Poa pratensis, Dichanthelium oligosanthes, Rumex acetosella* and *Aristida basiramea*, while the five most common species in mid plots were *P. pratensis, D. cognata, Ulmus minor, Ambrosia psilostachya* and *Koeleria macrantha*. Mean moisture (percentage water in soil) was 5.7% (± 1.0%, S.D.) in early plots and 6.1% (± 1.5%) in mid plots, while mean light availability (percentage of light available through canopy) was 56.2% (± 11.3%) in early plots and 52.4% (± 11.8%) in mid plots. Four of the study species (*A. millefolium*, *A. syriaca*, *A, tuberosa*, *S. scoparium*) were present at very low levels in the plots prior to experiment establishment (all <1% mean abundance). The other 12 species were present elsewhere in the field or surrounding fields but absent from all experimental plots.

On 15 June 2022, we planted 1,548 seedlings (approximately one month old) across 288 unmanipulated plots, with all existing vegetation undisturbed and intact. Plots were 1 m x 1 m, separated by 0.5 m walkways. Seedlings were planted into the central 0.5 m^2^ of each plot to limit possible edge effects (Fig. S3). Each plot received up to six individuals of a single species (some species had fewer than six individuals per plot due to low germination success, Table 1). We had three experimental enemy removal treatments (described below), and each species had three replicate plots per treatment. We therefore had 144 plots per successional stage (16 species × 3 treatments × 3 replicates), with species and treatments randomly assigned to plots to avoid spatial biases (Fig. S3). Before planting, we measured the height of all seedlings. Seedlings were watered inmediately after planting and as required throughout the first three weeks of the experiment to avoid excess mortality from drought stress.

The planted seedlings in each plot were assigned one of three enemy removal treatments: insecticide, insecticide+fungicide, or control. Pesticides were hand-painted on individual seedlings (i.e. only the seedlings were subject to enemy removal, not the whole community), using a foam brush and/or fine paintbrush until the leaves were visibly wet but not dripping. We did not do a fungicide-only treatment due to constraints on the overall area of the experiment and the time-intensive nature of treatment application; the effect of fungicide is interpreted as the difference between the insecticide and insecticide+fungicide treatments. We prioritised the insecticide single treatment as insect herbivory has generally been better studied (e.g. Turcotte *et al*. 2014), increasing comparability of our results. The insecticide was Marathon II (OHP, Inc., Mainland, PA, USA; 21.4% Imidacloprid) and the fungicide was Quilt (Syngenta Crop Protection, Inc., Greensboro, NC, USA; 7.5% Azoxystrobin and 12.5% Propiconazole). These pesticides have been shown not to affect plant growth (Seabloom *et al*., 2017, 2018). To control for the effect of painting, we painted water on plants in the control treatment. The treatments were applied every two weeks until the end of the growing season (31 August), with the first application occurring the day before being planted in the field (14 June; 7 applications across the season).

### 2.3. Data collection

Survival was assessed at approximately weekly intervals throughout the growing season, with seedlings being classed as alive if photosynthetic (green) tissue remained, and dead otherwise. We assessed survival at seven dates across eight weeks: 22 June, 30 June, 6 July, 18 July, 25 July, 31 July and 8 August. We recorded 10,836 individual observations in total (1,548 seedlings × 7 observation dates).

At the height of the growing season (beginning of August), we visually estimated percent cover (to the nearest %) of all species in the central 0.5 m^2^ of each plot, as well as bare soil, litter and disturbances (e.g. gopher mounds). There were 60 species (including our planted species) across all plots. We also visually estimated insect damage and fungal damage on our (surviving) focal seedlings to the nearest percent, taking the average of five haphazardly selected leaves on each plant. Tissue removal (chewing damage) from the margin or interior of leaves as well as leaf miner tracks were counted as insect damage, while lesions, rust spots and mildew were counted as fungal damage. We used the training program and techniques described in Xirocostas *et al*. (2022) to ensure accuracy in our estimates. All visual estimation for percent cover and damage was carried out by a single observer (J.I.B.).

On 15 August we measured % soil moisture content in all 288 plots, taking the average of four measurements in each plot using a Delta-T Devices HH2 Moisture Meter and SM300 Soil Moisture Sensor. On 16 August we measured light availability in each plot by taking the average of three light readings below the canopy with a Decagon Sunfleck PAR Ceptometer, dividing this by the light level above the vegetation layer and multiplying by 100.

We collected information on species phylogeny and functional traits to inform our statistical analyses. We built two phylogenetic trees using the V.PhyloMaker package (Jin & Qian, 2019). The first tree consisted of our planted 16 species, while the second consisted of all 60 species observed across all plots. We used the second tree to take a ‘focal-species’ approach (following Pinto-Ledezma *et al*., 2020), where we calculated the mean phylogenetic relatedness between our planted seedlings and all other species in the plot, weighted by the relative abundance of those species (abundance-weighted Mean Phylogenetic Distance, hereon ‘phylogenetic distance’). We collated data for five leaf traits [specific leaf area (SLA; mm^2^ mg^-1^), leaf dry matter content (LDMC, mg g^-1^), leaf area (mm^2^), leaf N (%) and leaf P (%)] for each of the 16 species. Sources of trait data were (Reich & Oleksyn, 2004; Craine *et al*., 2012; Catford *et al*., 2019) (see Methods S1 for full details). We selected these leaf traits to represent the broad axis of the plant fast-slow growth continuum (Craven *et al*., 2018).

Although our experiment used species native to the study region (Table 1), we examined whether they are naturalised in other parts of the world. We searched the GloNAF database (van Kleunen *et al*., 2019) for each of our 16 species, and recorded the number of regions in which they are reported to be naturalised (i.e. introduced by humans and have self-sustaining populations outside of their native range). The regions recorded in GloNAF are ‘basic recording units’ (i.e. TDWG level 4) where possible, and are recorded at the country level (TDWG level 3) otherwise. They range in size from 0.03 to 2,486,952 km^2^. Interpretations should thus be treated with caution as they encompass a broad range of sizes and levels of study effort (van Kleunen *et al*., 2015).

### 2.4. Statistical analysis

All statistical analyses were carried out in R v4.1.1 (R Core Team, 2021). See Supporting Information Code S1.

#### 2.4.1. Models explaining seedling survival

After confirming that pesticides reduced enemy damage (Fig. S4), we ran a series of models to explore factors influencing seedling survival. First, we used the proportion of seedlings surviving in a given plot as the response variable (model set 1a, Methods S2), and ran seven models (one for each observation date). Given high overdispersion in the data, we fitted generalised linear models using the quasibinomial family (logit link). Explanatory variables were treatment (control, insecticide, insecticide+fungicide), successional stage (early, mid), focal species, plot richness, phylogenetic distance, moisture, and light availability. We tested all two-way interactions: none were statistically significant (α=0.05). We therefore present results from models that only contain main effects. Statistical significance of a variable is determined by a χ^2^-based likelihood ratio test comparing models with and without the factor of interest. Diagnostics for these models (and the ones described below) models were inspected using the DHARMa package (Hartig, 2022) and found to be acceptable. Outliers for all of these and below models were identified using Cook’s distance. We ran models for each of the seven observation dates separately (following Charles *et al*., 2018), as the dates are not independent and so date cannot be included as a factor (if a seedling is dead at time *t*, it is guaranteed to be dead at *t* + 1). This approach also allows us to explicitly test the strength of the treatment effect at each time period. Given this multiple testing approach, we draw conclusions at a conservative Bonferroni-corrected α=0.007 (0.05/7). However, we also explicitly tested for a treatment*time interaction by using the number of new dead seedlings per plot as a response variable, and running a generalised linear model (Poisson family, log link) with a treatment*time interaction and successional stage, species, plot richness, phylogenetic distance, moisture, and light availability as main effects (model set 1b, Methods S2).

As differences in the initial size of seedlings may have affected seedling survival, we ran a set of seven mixed effects models (one for each observation date) at the individual seedling level, with binary survival (yes/no) as the response variable (model set 2, Methods S2). Models used the binomial family (logit link) and had initial height, treatment and successional stage as fixed effects, with plot nested within species as a random effect to account for the plot-level variables of phylogenetic distance, moisture, light and plot richness (these could not be directly included in the model as fixed effects due to severe multicollinearity, as they are identical for all seedlings in a given plot). Initial height was measured when seedlings were transplanted from the glasshouse to the field experiment. We used the package glmmTMB (Brooks *et al*., 2017) for these analyses.

#### 2.4.2. Exploring differences among species

Despite the lack of interaction between species and treatment in model set 1a, we ran individual-level models for each species independently, to quantify the effect of the treatment on survival at the species level in the final time period (model set 3, Methods S2). Models used the binomial family (logit link) and had initial height, treatment and successional stage as fixed effects, with plot as a random effect to account for phylogenetic distance, moisture, light and richness. We observed pronounced heterogeneity in the responses (i.e. effect sizes) of individual species to pesticide treatments, and explored whether this could be explained by phylogeny, plant traits or damage rates.

Using the phylogenetic tree of our 16 planted species, we tested whether more closely related species had more similar effect sizes (where ‘effect size’ refers to the benefit a species received from enemy removal treatments relative to controls). We used the phylosignal package (Keck *et al*., 2016) to test the null hypothesis that effect sizes were randomly distributed in the phylogeny. The phylogenetic signal was measured using the C_mean_ index, and the null hypothesis was rejected if p < 0.05 (Keck *et al*., 2016). We then carried out a PCA on leaf traits and extracted the first two axes (Fig. S2a). We examined the relationship between these PCA axes and both insecticide and insecticide+fungicide effect sizes for each species using generalised linear models (effect sizes logged to meet assumptions), with functional group (grass or forb) as a random intercept. We also compared effect sizes against the single traits of SLA and leaf N, as two of the most important plant traits determining rates of individual- and community-level enemy damage (Cappelli *et al*., 2020). Finally, we compared insect and fungal damage on our control plants with the insecticide and insecticide+fungicide effect sizes using simple linear models (effect sizes logged), to test whether species that received more damage in controls had larger treatment effect sizes. For the above three sets of analyses, two species were outliers based on Cook’s distance (*M. fistulosa* for the insecticide+fungicide treatment and *P. virgatum* for the insecticide treatment). We repeated analyses with these outliers excluded, which did not change overall results.

#### 2.4.3. Linking the effect of enemy release with naturalisation

We tested whether there was a relationship between species’ effect sizes (the benefit they received from treatments relative to controls) and the number of regions they are reported to be naturalised in. We used standard linear models with effect sizes and naturalised regions logged to meet the assumption of constant variance. The effect size (insecticide or insecticide+fungicide) was the independent variable, with number of naturalised regions globally as the dependent variable. We ran three alternative models for each effect size: forbs and grasses together, as well as separately. One species was a clear outlier based on Cook’s distance (*A. gerardii*: high effect sizes with zero naturalised regions); we repeated analyses with this outlier excluded, which did not change overall results.

## 3. Results

### 3.1. Factors affecting seedling survival

Pesticide application significantly reduced insect damage in the insecticide treatment, and reduced both insect and fungal damage in the insecticide+fungicide treatment (Fig. S4). Pesticide treatments significantly enhanced survival, in a way that depended on time (Fig. 1a; Tables S1 – S8). We present the plot-level models (model set 1a, based on 3–6 seedlings/plot), though we note the individual-level models (model set 2, based on individual seedlings) yielded identical results (Fig. S5, Tables S9 – S15). At the start of the experiment, there were no differences between control and pesticide-treated seedlings (χ^2^_2_=1.429, p=0.489); eight weeks after the experiment was established, survival of treated seedlings was 1.8 times higher on average than the control seedlings (χ^2^_2_=13.26, p=0.001; Fig. 1b). This difference was statistically significant at a Bonferroni-corrected α=0.007 for the last two sampling points (Tables S1 – S7). However, there was no difference between the insecticide and insecticide+fungicide treatments (Fig. 1). The Poisson model (model set 1b) confirmed the interaction between time and treatment, with the number of dead seedlings accumulating faster in control plots than treatment plots (Fig. S6, Table S8).

**Figure 1:**
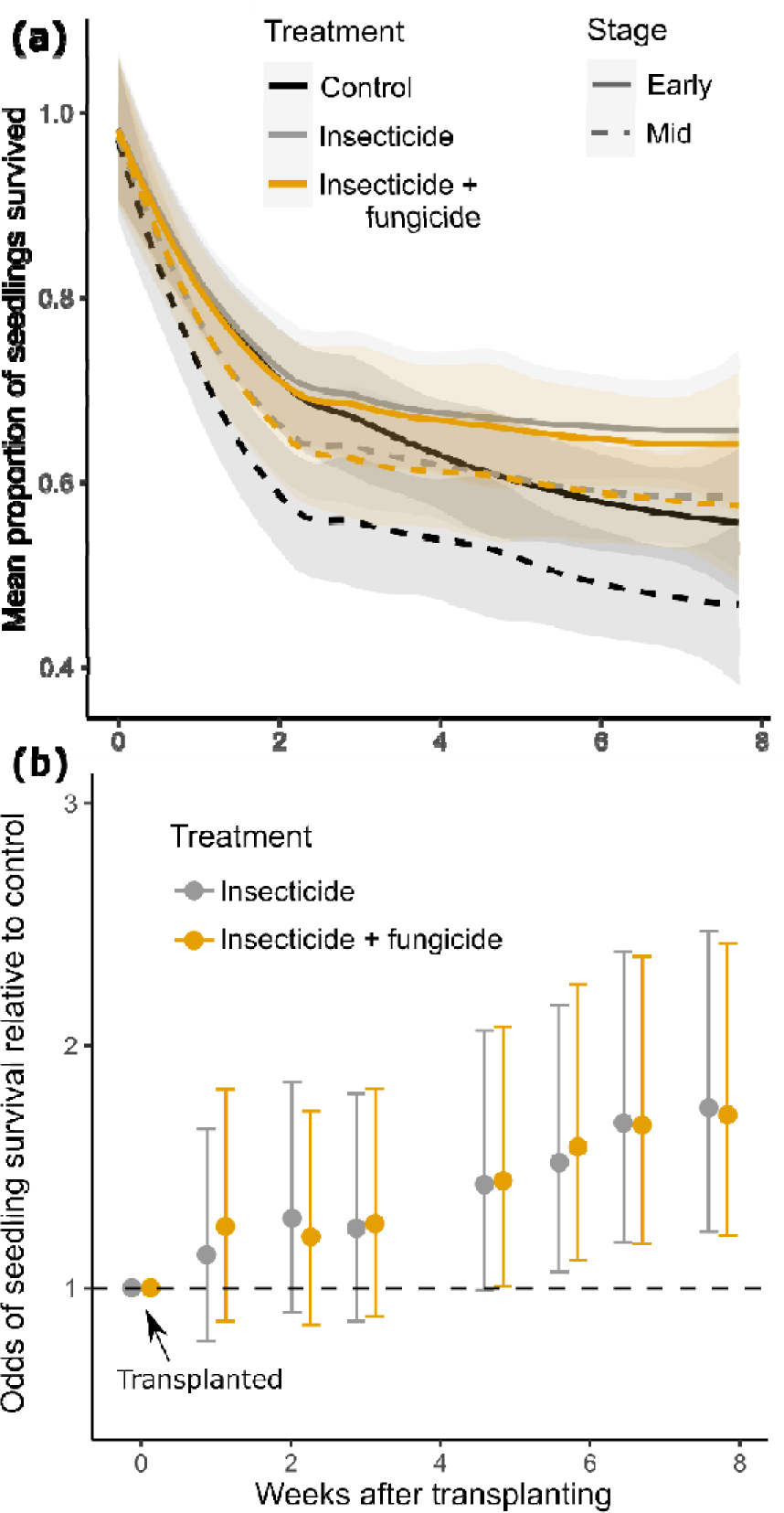
Pesticides enhanced seedling survival. (a) The overall raw proportion of seedlings per plot that survived, separated by treatment and successional stage. Lines are smoothed loess fits ± 95% C.I. (b) Effect size of treatments at each time point, averaged across all species (± 95% C.I.). Survival is measured as the proportion of seedlings in a plot surviving. The dashed horizontal line represents survival rates equal to the control group, averaged across all other factors.

Community successional stage, light availability and species richness in plots were also correlated with seedling survival. Based on the model at the final time point (Table S7), the odds of survival at the mid successional stage were only 0.51 times those at the early successional stage (χ^2^_1_=15.87, p<0.001; Figs. 1a, S7a). Survival of seedlings increased with light availability, a marginally significant effect (χ^2^_1_=2.767, p=0.096; Figs. 2a, S7b), and with plot species richness (χ^2^ =9.787, p=0.002; Figs. 2b, S7c). These effects were consistent in magnitude and statistical significance throughout the growing season (Fig. S7, Tables S1 – S7). Combined with the increasing influence of treatment, our ability to explain variation in survival therefore increased throughout the growing season, with the marginal R^2^ rising from 0.332 to 0.382 after eight weeks. Seedling survival was not related to seedling-community phylogenetic distance or plot soil moisture at any time during the growing season (Tables S1 – S8).

**Figure 2:**
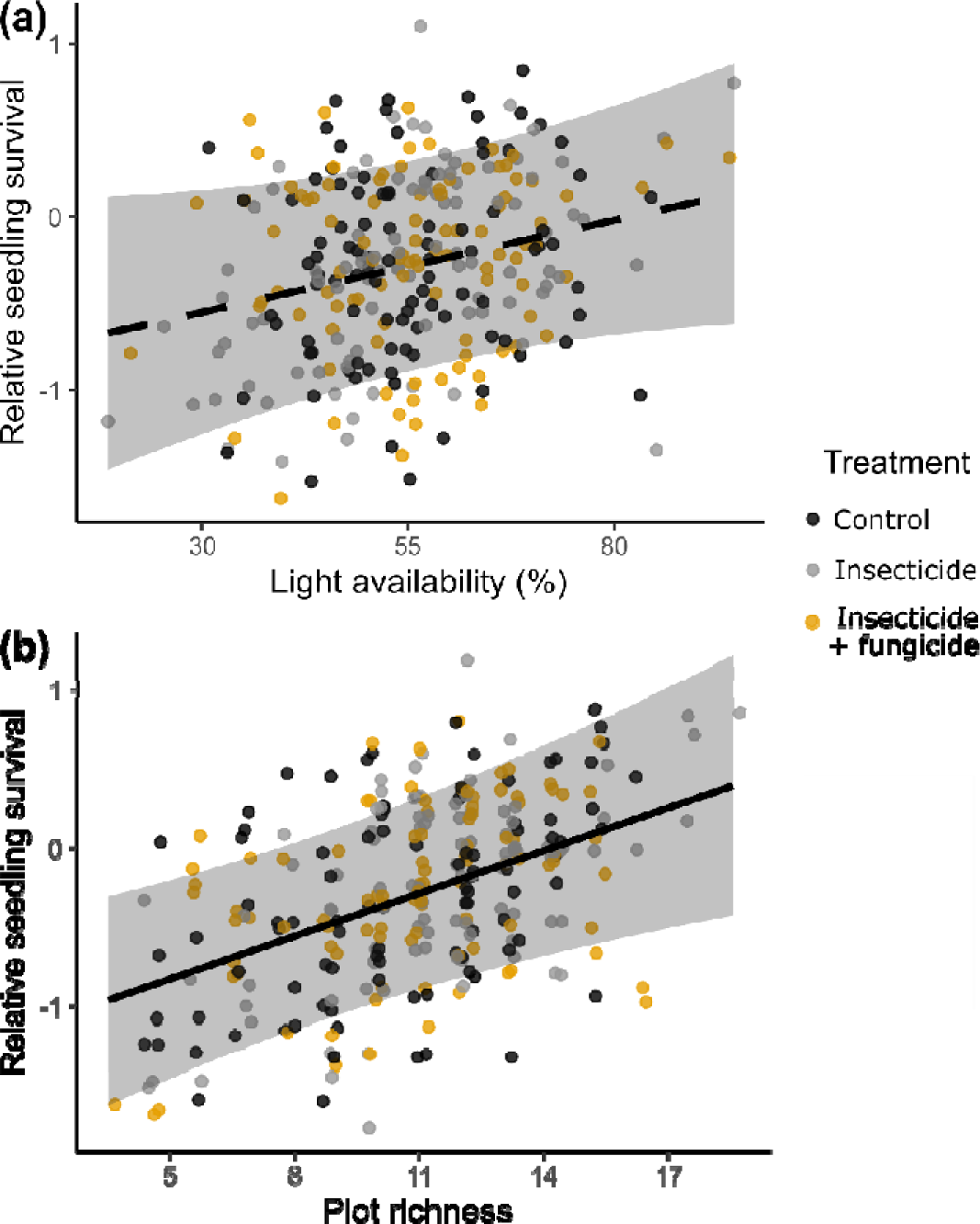
The effects of (a) light availability and (b) plot richness (total number of species in the plot) on the survival of seedlings. These are marginal effects plots from the final time point (8 August), holding all other factors constant. Dashed line indicates marginal significance.

### 3.2. Differences in response between individual species

There was high variation in the effect of treatments among species (Fig. 3; Tables S16 – S31). Our models found no statistically significant interaction between species and treatment (χ^2^ =25.43, p=0.704), suggesting the treatments were equally effective for all species. However, high stochasticity may have prevented detection of a statistically significant effect (Fig. 3), so we nevertheless explored whether species-specific differences in effect size could be explained by phylogenetic relatedness among species, species traits or rates of damage by insects or fungi. Two species (*M. fistulosa* and *P. virgatum*) had very large effect sizes for one of the treatments (Figs. 3, 4); trends remained the same after removing these outliers and repeating the analyses.

**Figure 3:**
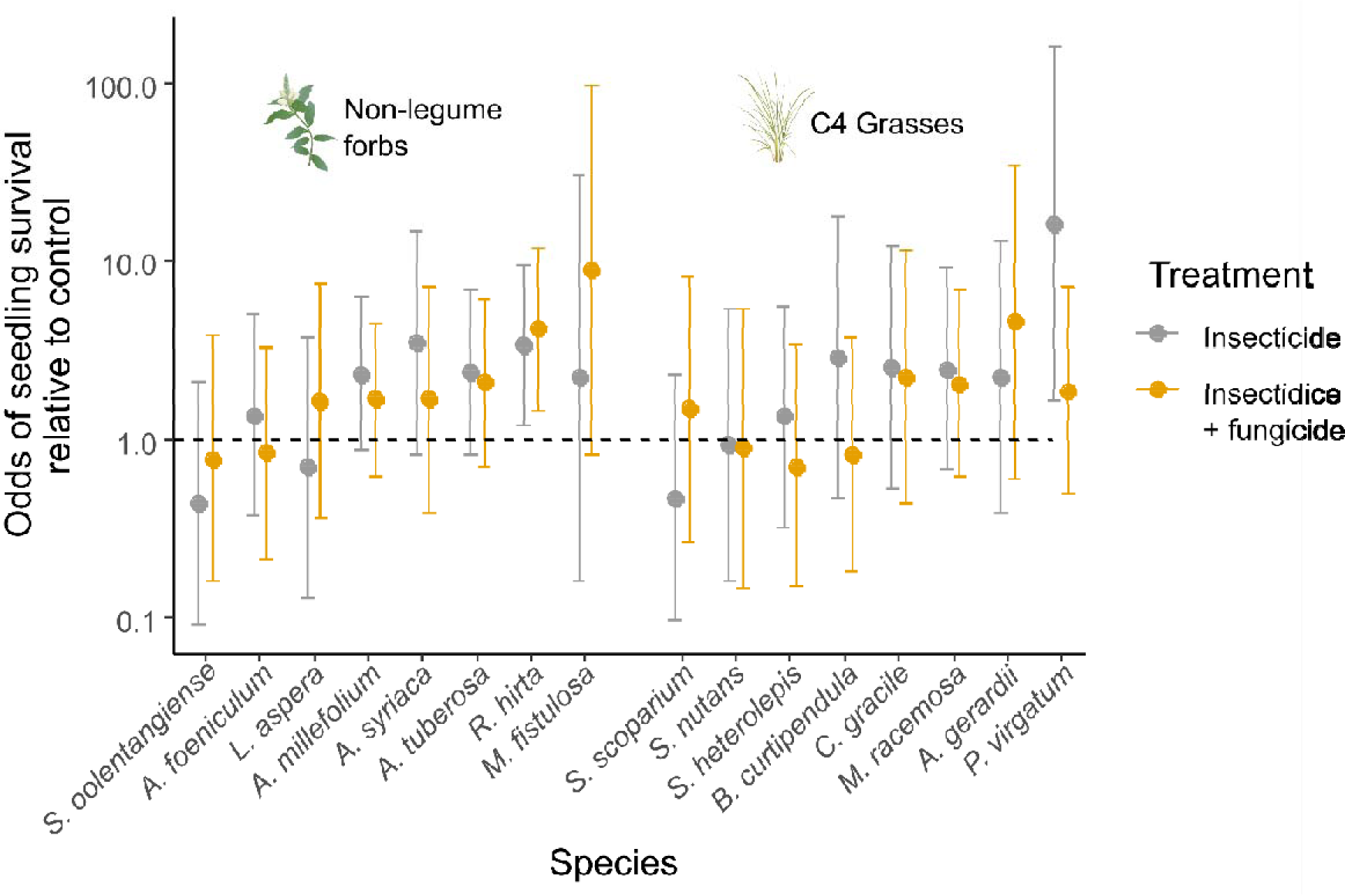
The effect sizes of insecticide and insecticide + fungicide treatments relative to the control after 8 weeks, separated by functional group and species. The dashed horizontal line represents survival rates equal to the control group, averaged across all other factors. Log scale on y-axis.

**Figure 4:**
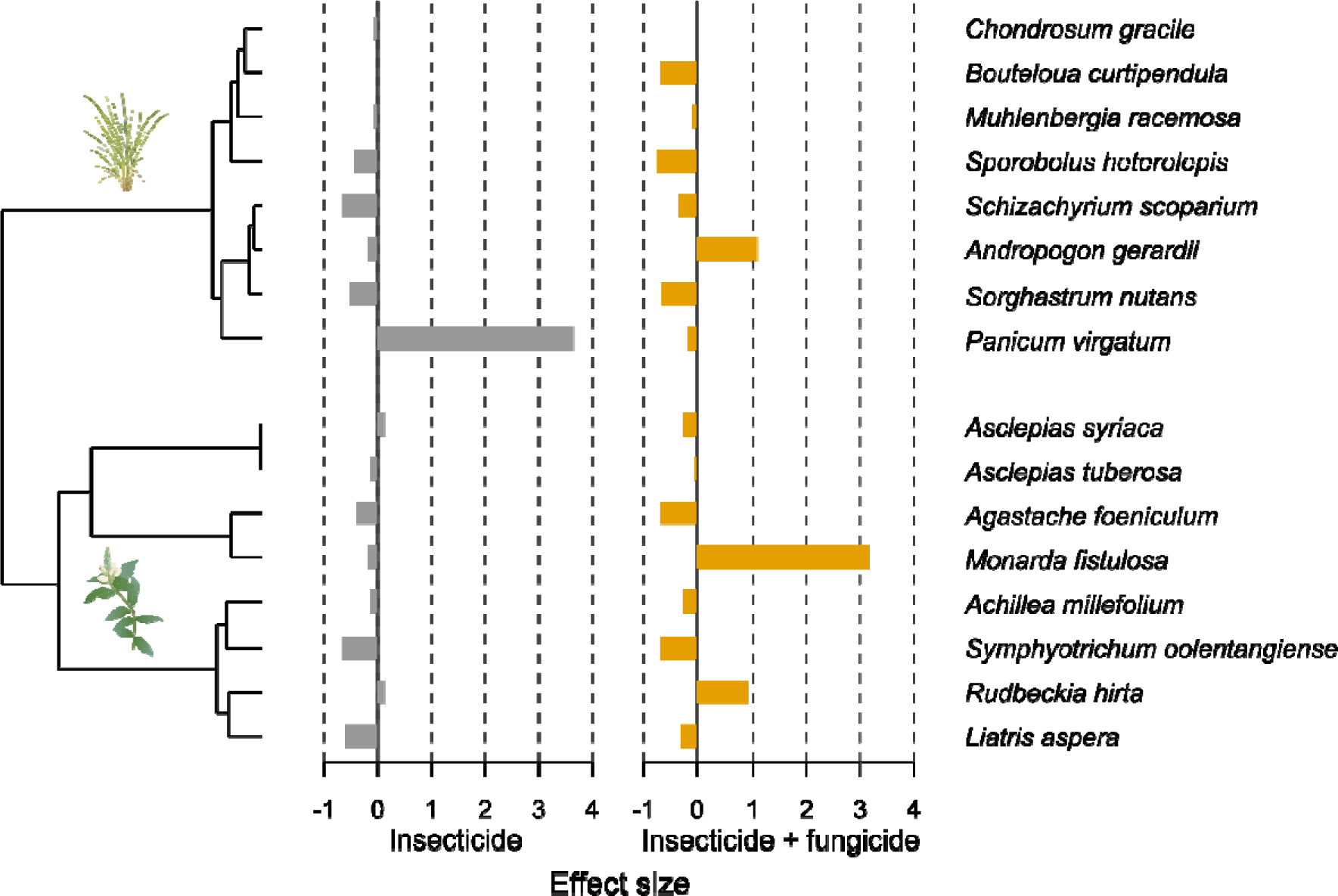
No statistically significant relationship between phylogeny of planted species and the effect size of insecticide treatment (left) and insectide+fungicide treatment (right). Note that effect size was scaled to have mean 0 in this analysis. Grasses are positioned above forbs.

Effect sizes were not related to phylogeny, for either the insecticide (p=0.782) or insecticide+fungicide treatment (p=0.925) (Fig. 4). Effect sizes were also unrelated to levels of damage on control plants (Fig. S8, Table S32). However, aboveground leaf traits (Fig. 5a) were correlated with effect sizes, with more resource-acquisitive plants showing higher benefit from the insecticide treatment in both functional groups (χ^2^_1_=4.049, p=0.044; Fig. 5b, Table S33). This relationship was also present but non-significant (α = 0.05) for the insecticide+fungicide treatment. Neither leaf SLA nor leaf N were significantly correlated with effect sizes, but they showed the same trend, with high-SLA and high-leaf N species showing larger positive effect sizes in both the insecticide and insecticide+fungicide treatments (Table S33).

**Figure 5:**
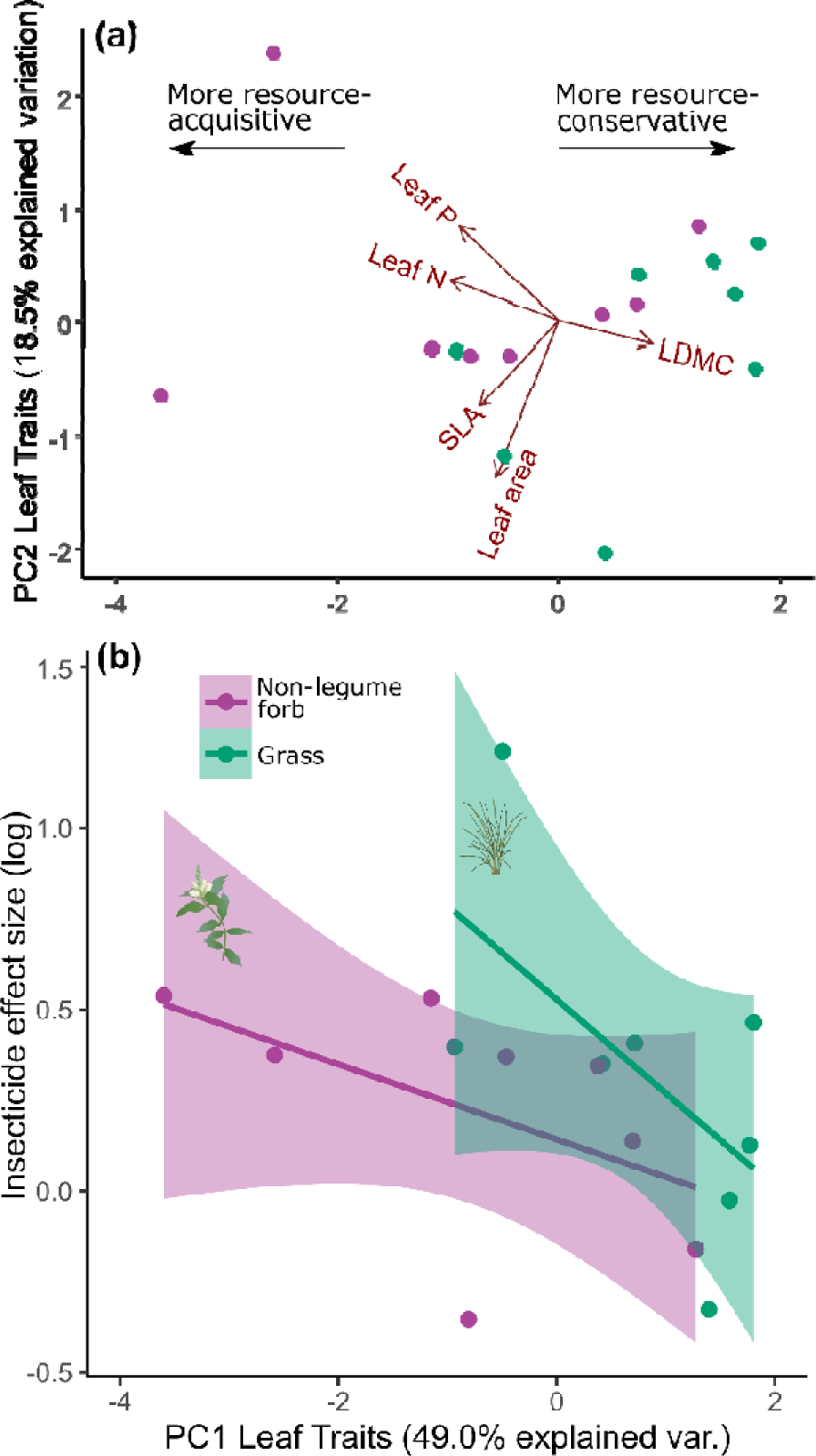
Relationship between leaf trait principal component analysis and effect sizes. Points show individual species and are coloured by functional group. (a) The first two PC axes with associated traits. (b) The further along PC1 (more resource-conservative) a species was, the lower the benefit from insecticide treatment. For full PCA results including species labels see Fig. S2.

Seedlings of different species showed different base survival rates throughout the season (χ^2^ =225.0, p<0.001; Fig. S9). We do not draw specific inferences from this pattern, as it could reflect different species-specific tolerances to transplantation or slight differences in time to germination rather than an ecological effect per se.

### 3.3. Relationship with naturalisation in invaded ranges

We found a strong positive relationship between treatment effect sizes and the number of regions where our experimental species are known to have naturalised (Fig. 6). The greater the benefit our species received from experimental enemy release, the more regions in which they are reported as being naturalised. This positive relationship was especially marked for the insecticide treatment (F_1,13_=8.613, p=0.012; Fig. 6a) but was also present (albeit statistically non-significant) for the insecticide+fungicide treatment (F_1,13_=2.861, p=0.117; Fig. 6b). This trend was also observed for forbs and grasses separately (Fig. 6c-f; Table S34). These results (both sign and statistical support) were identical when the outlier (*A. gerardii*) was included (Fig. S10, Table S34).

**Figure 6:**
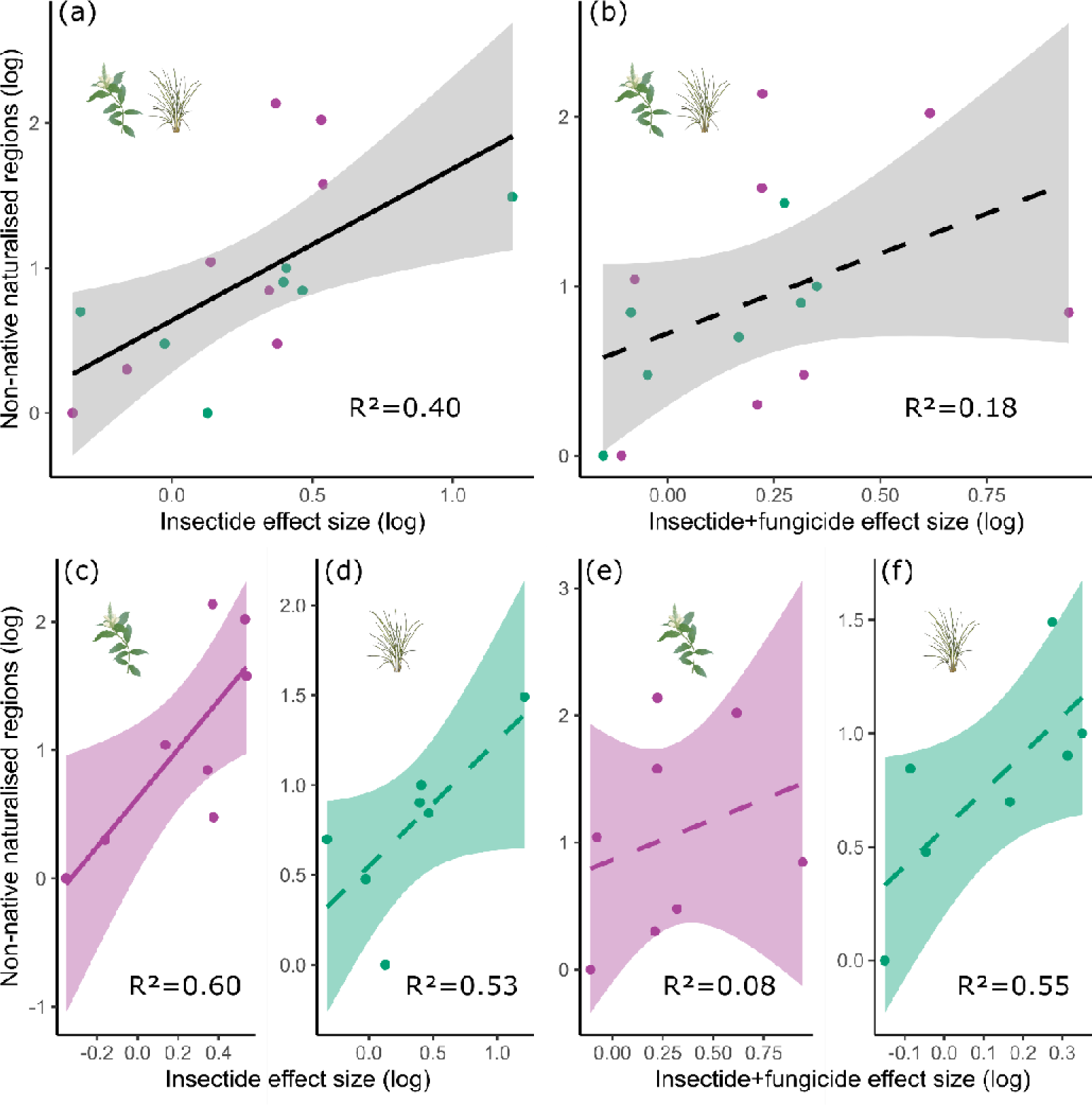
Relationship between treatment effect sizes (benefit of enemy release on seedling survival relative to controls) and the number of regions globally in which each species is reported as being naturalised (from the GloNAF database, van Kleunen *et al*., 2019). Relationships are shown for insecticide effect sizes (a, c, d) and insecticide+fungicide effect sizes (b, e, f), for both forbs and grasses combined (a, b) as well as for forbs separately (c, e) and grasses separately (d, f). Each dot is an individual species. Solid lines indicate statistical significance, dashed lines indicate statistical non-significance (though note high R^2^ values for several non-significant relationships, suggesting p values are related to low sample size rather than relationship strength).

## 4. Discussion

By hand-painting individual seedlings with pesticide to enforce enemy release, we found that enemy removal increased seedling survival of both non-legume forbs and C4 grasses in this grassland field experiment. Our result causally links the mechanism of enemy release with enhanced performance and considers survival in recruitment, two key gaps in the current enemy release literature. As we focused on 16 different species from two functional groups, we also demonstrate the wide scope of this result, emphasising the potential importance of enemy release in the colonisation phase of invasion, regardless of specific invader identity or functional group.

### 4.1. Enemy release increases early survival and recruitment of colonists

Pesticide application enhanced survival of our planted seedlings relative to controls, by an average of 80%. This effect was broadly consistent across successional stages and functional groups. Effect sizes for insecticide and insecticide+fungicide treatments were the same (Fig. 1): this may suggest that seedling survival is strongly reduced by insect herbivory but not fungal infection in this system. Alternatively, it could suggest that fungal release provides no additional benefit if seedlings have already been released from insects. Based on our experimental design with no separate fungicide treatment we cannot rule out the possibility that this would also be true for fungi; in other words, it could be that release from insects would provide no additional benefit if they have already been released from fungi.

Our results emphasise the importance of considering the impact of enemy release across the whole life cycle of exotic plants. It has been argued that only vertebrate herbivores prevent establishment of exotic plants as they consume whole plants or parts of plants, and that non-vertebrate enemies only limit invader abundance or spread once populations are established (Levine *et al*., 2004; Scurr *et al*., 2008; Zhang *et al*., 2018). Our findings suggest that, by removing tissue, insect herbivores may have a similar impact on seedlings as vertebrate herbivores, as even small levels of tissue removal will have a disproportionately large impact on plants that may only have a few leaves. The impact of fungal pathogens, which degrade but do not actively remove tissue, may not reduce plant survival but instead limit biomass and competitive ability later in life (Mitchell, 2003; Zaret *et al*., 2022). Literature suggests that the effect of enemy release may be observed for certain life stages but not others: for example, pesticides applied to a whole forest community reduced invader recruitment success (due to disproportionately benefiting existing native vegetation), but later improved the height and diameter of invading trees (Heckman *et al*., 2022). Therefore, considering various life stages is important in the context of enemy release. Grass and forb invaders may significantly benefit from release in terms of seedling survival, even if previous studies suggest it makes little difference to the competitive ability of adults (e.g. Dong *et al*., 2018; Katz & Ibáñez, 2017).

The survival benefit of release from insect herbivores was equally observed in both C4 grasses and non-legume forbs (Fig. 3). The benefit to C4 grasses was somewhat unexpected, given previous work has found little benefit of pesticides on C4 grasses relative to forbs (Seabloom *et al*., 2018). However, this previous work focused on biomass rather than early survival, again emphasising the importance of considering different exotic life-history stages. While in general fungal pathogens are more common on grasses and insect herbivores are more common on forbs (Ebeling *et al*., 2022; Fricke *et al*., 2022), a pattern also borne out in our experiment (Fig. S8), it appears that both forbs and C4 grasses can benefit from release of insect herbivory during recruitment. C4 grasses may be particularly vulnerable as seedlings relative to adults (Dybzinski & Tilman, 2012), and thus also suffer reduced survival from insect tissue removal.

Within each plant functional group, we observed variation in treatment effect size that was linked to aboveground traits, with resource-acquisitive plants benefiting more from enemy release (Fig. 5). The absence of a clear phylogenetic signal (Fig. 4) affirms the need to incorporate information on functional traits into predictions of invasion success, as phylogeny alone may not always predict meaningful ecological differences (Cadotte *et al*., 2017). Resource-acquisitive species typically invest in fewer defences and are more vulnerable to enemies (Grutters *et al*., 2017; Heckman *et al*., 2019; Cappelli *et al*., 2020). Resource-acquisitive plants may therefore be especially damaged and should receive greater benefits from enemy release (Blumenthal *et al*., 2009). Resource-conservative plants should benefit little from enemy release as they are heavily defended so less susceptible to damage. However, on average, resource-conservative species still benefited in our experiment. As juvenile plants have lower capacities for defence than adult plants (Bruns *et al*., 2022), the hypothesised relationship between plant strategies and benefits of enemy release may be more decoupled during plant recruitment. Enemy release could therefore benefit the initial survival of invaders regardless of species traits, though the leaf economics spectrum highlights where release during recruitment may be particularly beneficial (Fig. 5b). This trend does not seem to fit with our observation that the benefits of enemy release were unrelated to damage on control plants (Fig. S8): we would expect that more highly damaged plants in the controls should benefit the most from enemy release. We suggest this lack of relationship is because we only assessed damage once, at the end of the season. Highly damaged plants may have already died, thus lowering average damage on (surviving) control seedlings and masking the expected relationship between damage and enemy removal benefits.

Successional stage, light and plot-level richness were also important in determining survival of our seedlings. Although soil nitrogen, not light, is the key limiting resource at Cedar Creek (Tilman, 1990), it is possible that the short heights of seedlings mean they still receive some benefit from increased light availability (Fig. 2a) (e.g. Johnson *et al*., 2020). Similarly, mid succession communities tend to be more species-rich and nutrient-poor than early succession communities, characteristics that increase resistance to invasion and colonisation (Fridley *et al*., 2007; Catford *et al*., 2020; Liu *et al*., 2021). Indeed, our mid plots had an average of two more species per plot than early plots and total summed plant cover was 20% higher in mid plots, potentially exerting stronger competitive pressure on seedlings (Fig. 1a). However, within each site we unexpectedly observed a slight increase in survival of pesticide-exposed seedlings as plot richness increased (Fig. 2b). This result has also been observed before, though in tree communities studied over longer time periods (Liu *et al*., 2022). We suggest that plot-level richness may be correlated with reductions in total enemy pressure, as increased local diversity has been shown to decrease infection prevalences on individual plants (Rottstock *et al*., 2014). While we observed no relationship between plot richness and aboveground damage rates in our study (Fig. S4), it is possible that increased richness reduced the impact of unmeasured belowground pathogens.

We found a positive relationship between the benefit that a species received from enemy release, and the number of regions globally in which that species is reported as being naturalised (Fig. 6). Once again, this was observed for both forbs and grasses separately (Fig. 6c-f). There are many reasons a species may successfully naturalise outside of their native range (Díaz *et al*., 2023), so this correlation cannot be taken as direct evidence that enemy release facilitated the spread of these species in other parts of the world. For example, as we also found a positive correlation between the effect of enemy release and leaf-acquisitive traits (Fig. 5b), the degree of naturalisation could be related to a ‘fast’ life-history strategy (van Kleunen *et al*., 2010b), independent of enemy release. Further caution is required, as regions used in GloNAF are not necessarily equivalent to ecoregions, and being successful in multiple different biomes is different to being successful in multiple sites within the same biome. Nevertheless, the positive correlations between naturalisation, life history strategy and enemy release provide a compelling hypothesis that the extent of non-native naturalisation is driven by the degree to which a species can benefit from enemy release. Further research testing this hypothesis would be valuable.

### 4.2. Implications for experimental tests of the ERH

Many experimental studies of the ERH in natural communities are indirect (e.g. Heckman *et al*., 2017; Suwa & Louda, 2012). They rely on naturally occurring exotic invaders (which may be experiencing different levels of enemy release, or none at all) and plot-level pesticide applications, leaving them unable to disentangle interactions between community diversity and biomass, enemy presence, and invader performance (Fig. S1). In contrast, we enforced enemy release by applying pesticides to individual plants and are thus better able to examine causal links between the key mechanism of enemy release and seedling survival. Enemy release is necessarily species-specific, yet people typically investigate it in the field by releasing an entire community from enemies. We strongly recommend that future ERH experiments that use pesticides focus on individual-level applications, to directly link enemy release with metrics of plant performance.

We included two successional stages in our field experiment. Despite being unreplicated, the resident plant taxonomic and functional conditions of our two sites were reflective of nearby fields abandoned from agriculture at similar times (Catford *et al*., 2023) and, as such, were characteristic of sites at a similar successional stage. Successional stage was correlated with seedling survival, suggesting community composition is an important consideration in recruitment success (Fig. 1a), but the positive effect of enemy release was still observed in both types of communities. This trend aligns with previous work that found that effects of enemy release are consistent regardless of community structure (Suwa *et al*., 2010; Heckman *et al*., 2017). However, other studies suggest that the same plant species at sites with very different nutrient regimes may show different responses to enemy release (Dewalt *et al*., 2004; Jia *et al*., 2020). The generality of our results should be further tested by exploring survival in response to enemy release across different environments. More broadly, understanding how ecological context interacts with enemy release is vital to understanding the role of enemy release in global invasions (Brian & Catford, 2023).

As our aim was to test the effect of enemy release, not determine whether enemy release occurs “in nature”, we focused on species that were native to and extant in the study region (though very rare or absent in our specific plots). This was to ensure the presence of both specialist and generalist enemies, making it more likely that our pesticide applications would reduce damage rates and effectively simulate the key mechanism of enemy release (Fig. S4). The correlation between our results on our 16 target species and the extent to which these species are naturalised in other parts of the world suggests that our approach holds relevance for investigating possible causes of invasion success, though we acknowledge that our plants were adapted to the local conditions and constrained within the trait space of plants that normally occur in the region. While we selected plants from two functional groups and a broad range of trait space (Fig. S2), future work should test whether our results hold over a broader suite of species, including field manipulations of exotic seedlings where possible. Given that exotic species tend towards resource-acquisitive strategies (van Kleunen *et al*., 2010a, b) – characteristics we found to enhance the benefits of enemy release – it may be that our study on natives underestimates the benefits for exotic seedling survival, as our species may be less resource-acquisitive than the average exotic species.

### 4.3. Conclusion

We have demonstrated using a grassland experiment that the mechanism of enemy release can significantly enhance the survival of seedlings. We suggest that increasing early survival, by a mean of 80% across our 16 study species, could significantly enhance recruitment. The consistency of this result, being observed across C4 grass and non-legume forb seedlings and for seedlings transplanted into communities at different successional stages, suggests that enemy release-mediated recruitment may be an important but understudied aspect of plant species invasions. Even if enemies rapidly accumulate on invaders (Hawkes, 2007; Ivison *et al*., 2023), brief release during plant recruitment may be sufficient for invaders to gain a foothold in their new ranges.

## Acknowledgements

Funding for the project comes from the European Research Council (ERC) under the European Union’s Horizon 2020 research and innovation programme (grant agreement No. [101002987]). We are very grateful to Troy Mielke, Kally Worm and the rest of the 2022 Cedar Creek staff and interns for their enormous help in the establishment and maintenance of the experiment. Additional field assistance was provided by Zoe Sims and Laura Byrne. JIB is also very grateful for the advice, help and friendship throughout the field season from Maggie Anderson, Maria Park and Sydney Hedberg. Maggie Anderson took the drone photos in Fig. S3. Three anonymous reviewers provided valuable comments that improved a previous version of this manuscript.

## Author contributions

Jane A. Catford and Joshua I. Brian conceived the idea and designed the experiment. Joshua I. Brian established the experiment. Joshua I. Brian and Harry E. R. Shepherd collected data. Joshua I. Brian, Harry E. R. Shepherd and María Ángeles Pérez-Navarro analysed data. Joshua I. Brian drafted the manuscript, and all authors significantly contributed to revisions.

## Statement on inclusion

Our study includes authors with long-term collaborations in the country of interest and experience of working in the research site. Research staff at the site were consulted at all stages of experimental design, set-up and sampling.

## Data availability

For the purposes of peer review, all data and code supporting the analyses have been deposited in a public GitHub repository for inspection: https://github.com/brianjosh/enemyrelease If the paper is accepted, these data will be permanently accessioned using Dryad.

## Supporting Information

**Methods S1:** Acquisition and processing of species trait data

**Methods S2:** Description of model sets used in analysis

**Figure S1:** Issues with current community-level experiments of the enemy release hypothesis

**Figure S2:** The first two principal components for trait variation in leaf traits and belowground traits, with each species labelled

**Figure S3:** Experimental design

**Figure S4:** Effectiveness of pesticide treatment for reducing insect damage (insecticide treatment) and for reducing insect and fungal damage (insecticide+fungicide treatment)

**Figure S5:** Effects of pesticide treatment on individual seedling survival relative to control seedlings

**Figure S6:** The number of new dead seedlings per plot at each sampling event, split by treatment

**Figure S7:** The effect of successional stage, light availability, and species richness in determining the odds of survival of planted seedlings

**Figure S8:** Relationship between effect sizes for insecticide treatment, insecticide + fungicide treatment, and damage rates on control plants

**Figure S9:** Proportion of seedlings of each species surviving per plot, separated by successional stage and averaging across all other factors

**Figure S10:** Relationship between effect sizes and number of naturalised regions for each species, including the outlier A. gerardii

**Tables S1-S7:** Results from model set 1a (quasibinomial models exploring proportional seedling survival at the plot level at each time point)

**Table S8:** Results from model set 1b (poisson model exploring number of new dead seedlings through time)

**Tables S9-S15:** Results from model set 2 (binomial models exploring individual seedling survival at each time point)

**Tables S16-S31:** Results from model set 3 (binomial models exploring individual survival per species at the final time point)

**Table S32:** Relationship between damage on control plants and treatment effect sizes

**Table S33:** Relationship between plant traits and treatment effect sizes

**Table S34:** Relationship between effect sizes and number of non-naturalised regions that experimental species are reported in

