## Supporting Information for "Aboveground enemy release increases seedling survival in grasslands"

| Methods | | |
| --- | --- | --- |
|  | Methods S1: Acquisition and processing of traits | 2 |
|  | Methods S2: Model sets | 3 |
| Figures | | |
|  | Figures S1 – S10 | 5 |
| Tables | | |
|  | Tables S1 – S34 | 13 |

**Supporting Information: Methods**

**Methods S1: Acquisition and processing of traits**

For each of the 16 experimental species, we obtained five leaf traits: specific leaf area (SLA; mm^2^ mg^-1^), leaf dry matter content (LDMC, mg g^-1^), leaf area (mm^2^), leaf N (%) and leaf P (%). To demonstrate that our species cover a broad range of trait space, we also selected four root traits: specific root length (SRL, m g^-1^), root nitrogen (mg g^-1^), root diameter (mm) and root density (g cm^-3^).

Leaf traits were collected from measurements taken at the study site in neighbouring experiments (Catford et al., 2019). As trait data from Cedar Creek was insufficient for leaf N and P, we filled gaps firstly from a comparative North American grassland site (Craine et al., 2012) and then a global dataset (Reich & Oleksyn 2004).

Root traits were extracted from the GRooT database (Guerrero-Ramírez et al., 2021). As there was insufficient data for root trait values from Cedar Creek, we extracted trait values for both potted and field studies conducted in Minnesota and neighbouring states situated within the continental biome, the climatic biome of our study location. Trait values were then log-transformed and corrected for study design and source location using linear mixed-effects models as in Bergmann et al. (2020), with model residuals extracted for further use. In total, this provided 94% of species-level leaf traits and 92% of species-level root traits. Missing leaf and root trait values were then phylogenetically imputed using the R package *Rphylopars* (Goolsby et al., 2022).

*References*

Bergmann, J., Weigelt, A., van Der Plas, F., Laughlin, D. C., Kuyper, T. W., Guerrero-Ramirez, N., ... & Mommer, L. (2020). The fungal collaboration gradient dominates the root economics space in plants. *Science Advances*, **6**, eaba3756.

Catford, J. A., Smith, A. L., Wragg, P. D., Clark, A. T., Kosmala, M., Cavender‐Bares, J., ... & Tilman, D. (2019). Traits linked with species invasiveness and community invasibility vary with time, stage and indicator of invasion in a long‐term grassland experiment. *Ecology Letters*, **22**, 593-604.

Craine, J. M., Towne, E. G., Ocheltree, T. W., & Nippert, J. B. (2012). Community traitscape of foliar nitrogen isotopes reveals N availability patterns in a tallgrass prairie. *Plant and Soil*, **356**, 395-403.

Goolsby, E., Bruggeman, J., & Ane, C. (2022). Rphylopars: Phylogenetic Comparative Tools for Missing Data and Within-Species Variation. R package version 0.3.9, https://CRAN.R-project.org/package=Rphylopars.

Guerrero‐Ramírez, N. R., Mommer, L., Freschet, G. T., Iversen, C. M., McCormack, M. L., Kattge, J., ... & Weigelt, A. (2021). Global root traits (GRooT) database. *Global Ecology and Biogeography*, **30**, 25-37.

Reich, P. B., & Oleksyn, J. (2004). Global patterns of plant leaf N and P in relation to temperature and latitude. *Proceedings of the National Academy of Sciences*, **101**, 11001-11006.

**Methods S2: Model sets**

*Model Set 1: Seedling survival at the plot level.*

*1a: Quasibinomial models exploring proportional seedling survival at the plot level.*

*y_i_^t^* = Proportion of seedlings in a plot that survived at time *t* (sampling event 1 to 7), where *i* is a given plot (1 – 288).

E(*y_i_^t^*) = log(P[*y_i_^t^* = 1]/(1 - P[*y_i_^t^* = 1]) = Intercept + treatment + species + awMPD + successional stage + light + moisture + richness.

Var(*y_i_^t^*) = *φ*E(*y_i_^t^*)*(1 - E(*y_i_^t^*)), where *φ* = overdispersion parameter.

*1b: Poisson model exploring number of new dead per plot at each time point.*

*y_i_* = Number of new dead seedlings between time *t* and *t – 1*, where *i* is a given plot (1 – 288).

E(*y_i_*) = log(*y_i_*) = Intercept + treatment*time + species + awMPD + successional stage + light + moisture + richness.

Var(y*_i_*) = E(*y_i_*)

*Model Set 2: Binomial models exploring individual seedling survival.*

*y_i_^t^* = Seedling survival status (1 = yes, 0 = no) at period *t* (sampling event 1 to 7), where *i* is an individual seedling (1 – 1548).

E(*y_i_^t^*) = log(P[*y_i_^t^* = 1]/(1 - P[*y_i_^t^* = 1]) = Intercept + treatment + height + successional stage + (1|species/plot).

Var(*y_i_^t^*) = E(*y_i_^t^*)*(1 - E(*y_i_^t^*)).

*Model Set 3: Binomial models exploring individual survival per species at time 7 only.*

*y_i_^s^* = Seedling survival status (1 = yes, 0 = no) for seedlings of species *s* (species 1 - 16), where *i* is an individual seedling (1 – 1548).

E(*y_i_^s^*) = log(P[*y_i_^s^* = 1]/(1 - P[*y_i_^s^* = 1]) = Intercept + treatment + height + successional stage + (1|plot).

Var(*y_i_^s^*) = E(*y_i_^s^*)*(1 - E(*y_i_^s^*)).

**Supporting Information: Figures**

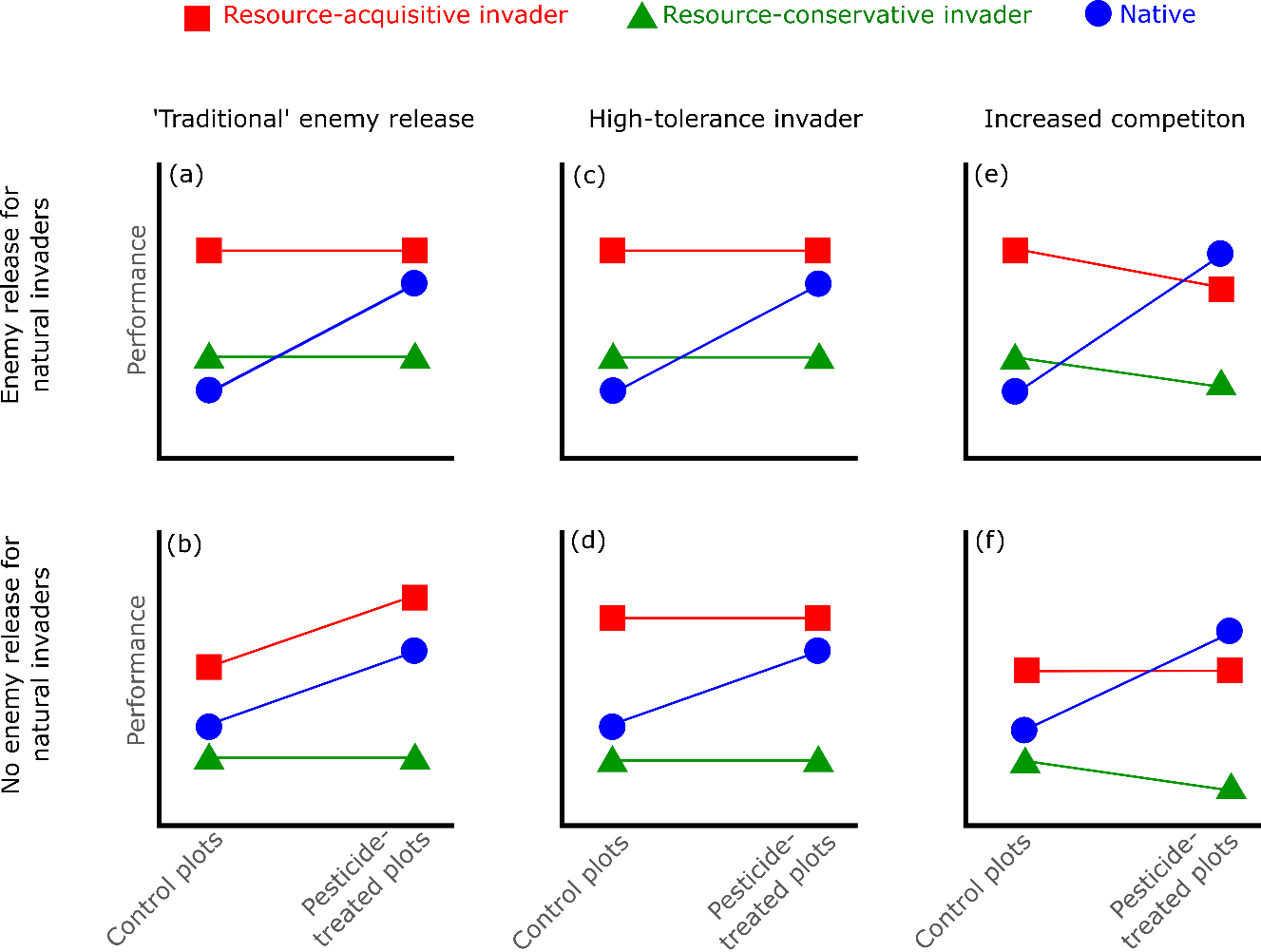

Figure S1: Issues with current community-level experiments of the enemy release hypothesis. Experiments typically apply pesticides to whole plots and examine the responses of naturally occurring invaders in the plots, compared to native species. The principle behind this approach is that invaders experiencing enemy release should not benefit from pesticide application, while native species do (a). If invaders are not experiencing enemy release, then they should also benefit from pesticide application (b). An initial problem with this approach is that this expectation does not hold for resource-conservative invaders: they typically have high defence levels and so experience similar success with or without enemies, and so should be largely unaffected by pesticide application regardless of whether they have experienced enemy release (identical green lines in (a) and (b)). Other factors also confound interpretations of this experimental design. For example, resource-acquisitive invaders can show high tolerance (recovery from enemy damage) through fast growth, and so have high performance even in the presence of enemies, and so pesticide application would make little difference, whether or not they have been released from enemies (compare (c) and (d)). Finally, whole-plot pesticide application can increase native biomass, and so might increase competition on invaders. Therefore, resource-acquisitive invaders already experiencing natural enemy release may decline in performance as they lose competitive advantage (e), while resource-acquisitive invaders that are not experiencing enemy release should simultaneously benefit from pesticide application and suffer from increased competition, and so appear to be unchanged (f). Finally, natural enemies accumulate through time: without clear knowledge of invader population age, it would be easy to conclude that the invaders are not benefiting from enemy release, but they may have benefited at the beginning of the invasion process. Note the difficulties in disentangling (a), (c), (d) and (f) for high-resource invaders, and (a) – (d) as well as (e) – (f) for low-resource invaders.

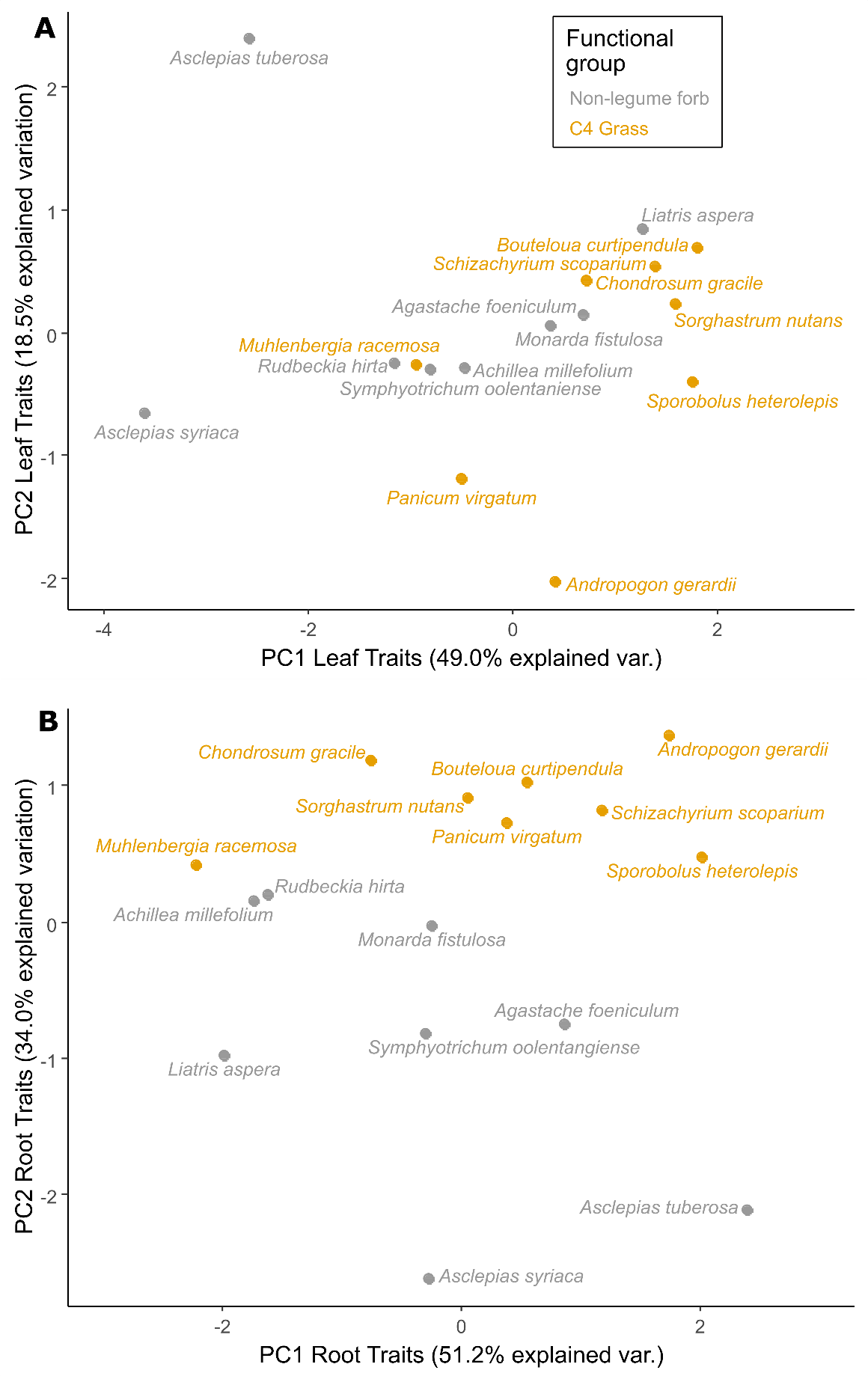

Figure S2: The first two principal components for trait variation in leaf traits (A) and belowground traits (B), with each species labelled.

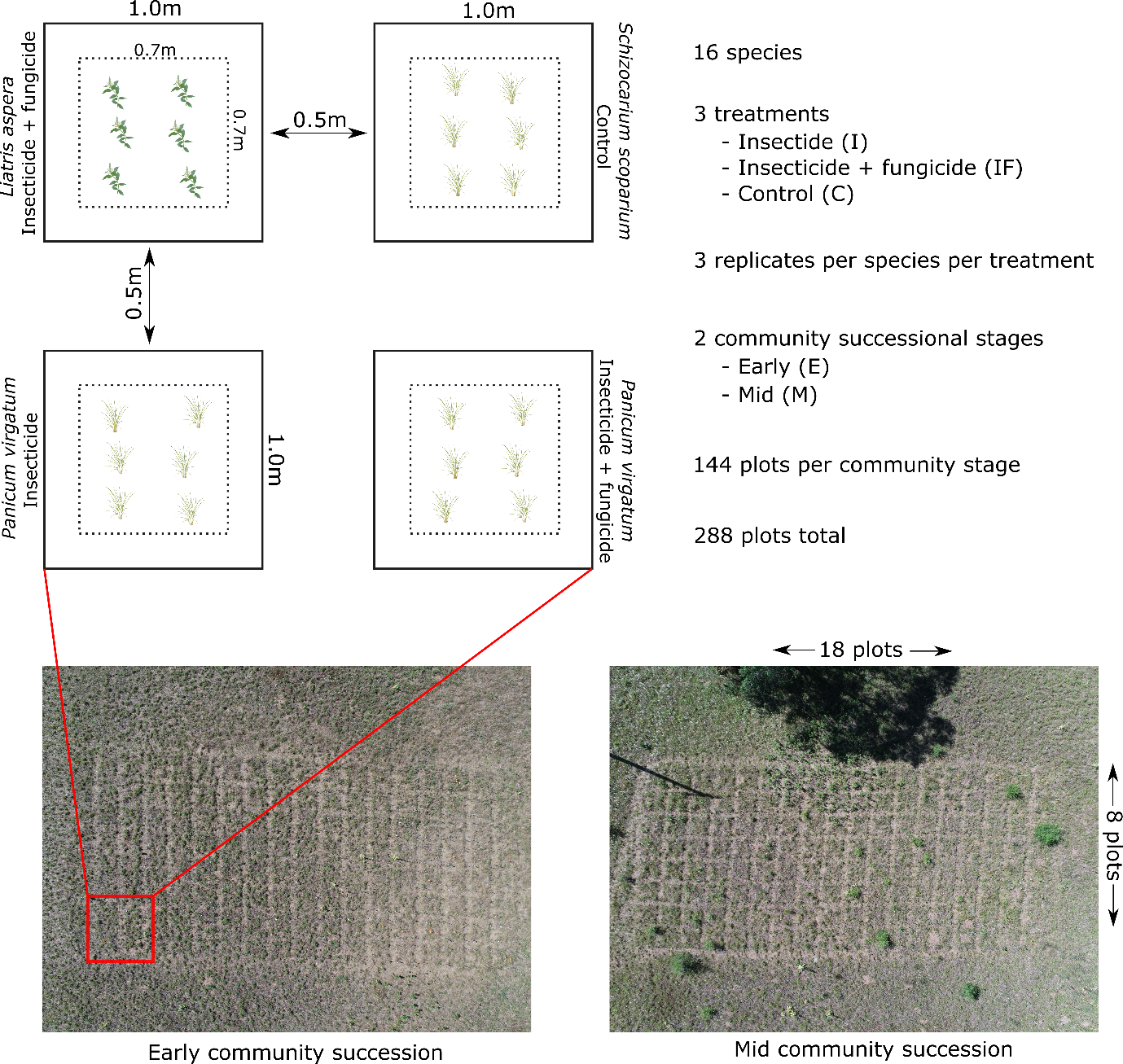

Figure S3: Design of the experiment, detailing plot set up, dimensions and replication. Species and treatment arrangement were completely randomised across the 144 plots at each community stage, but the arrangement was the same at both the early and mid community stages. The details of four plots are shown as an example. Drone photos taken by Maggie Anderson, University of Minnesota. Both photos taken 27^th^ July 2022.

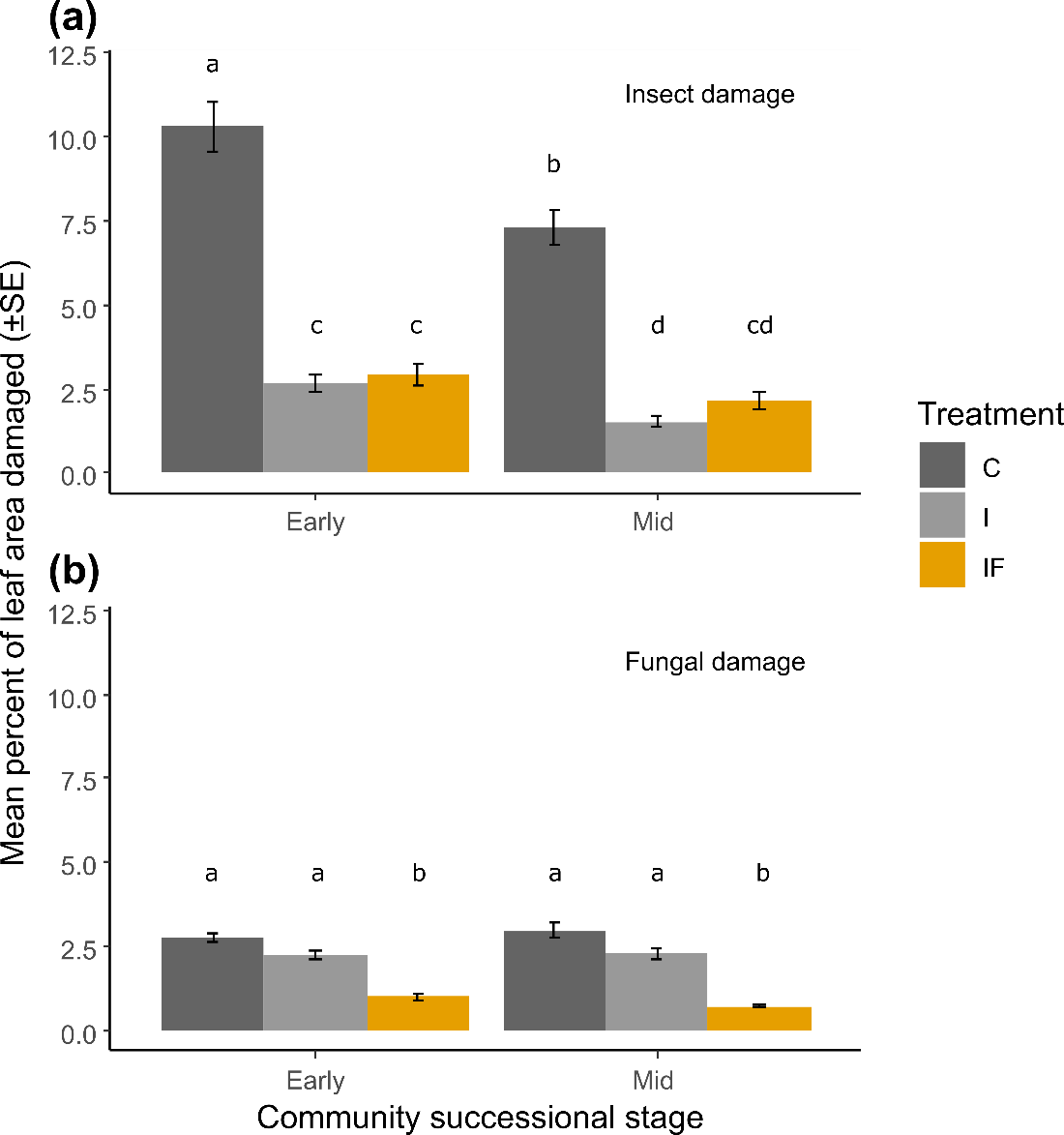

Figure S4: Mean leaf damage on planted seedlings by treatment (control, C; insecticide, I; insecticide + fungicide, IF), for (a) insects (chewing and leaf mining damage); and (b) fungi (lesions, rust spots and mildew). Letters above bars represent groupings as determined by post hoc pairwise Wilcoxon tests. Pesticide application significantly reduced damage on our treated seedlings (insect damage, χ^2^_2_=127.9, p<0.001; fungal damage: χ^2^_2_=84.2, p<0.001). Insect damage was on average 3.8 times lower in both the insecticide and insecticide+fungicide treatments relative to controls, while fungal damage was on average 3.1 times lower in the insecticide+fungicide treatment relative to both insecticide and control treatments. There was marginally less insect damage on seedlings in the mid succession communities relative to the early succession communities, but no differences in fungal damage between seedlings in the two community types. There was also no effect of plot richness on damage after accounting for all other factors (insect damage: χ^2^_1_=0.277, p=0.599; fungal damage: χ^2^_1_=0.603, p=0.438).

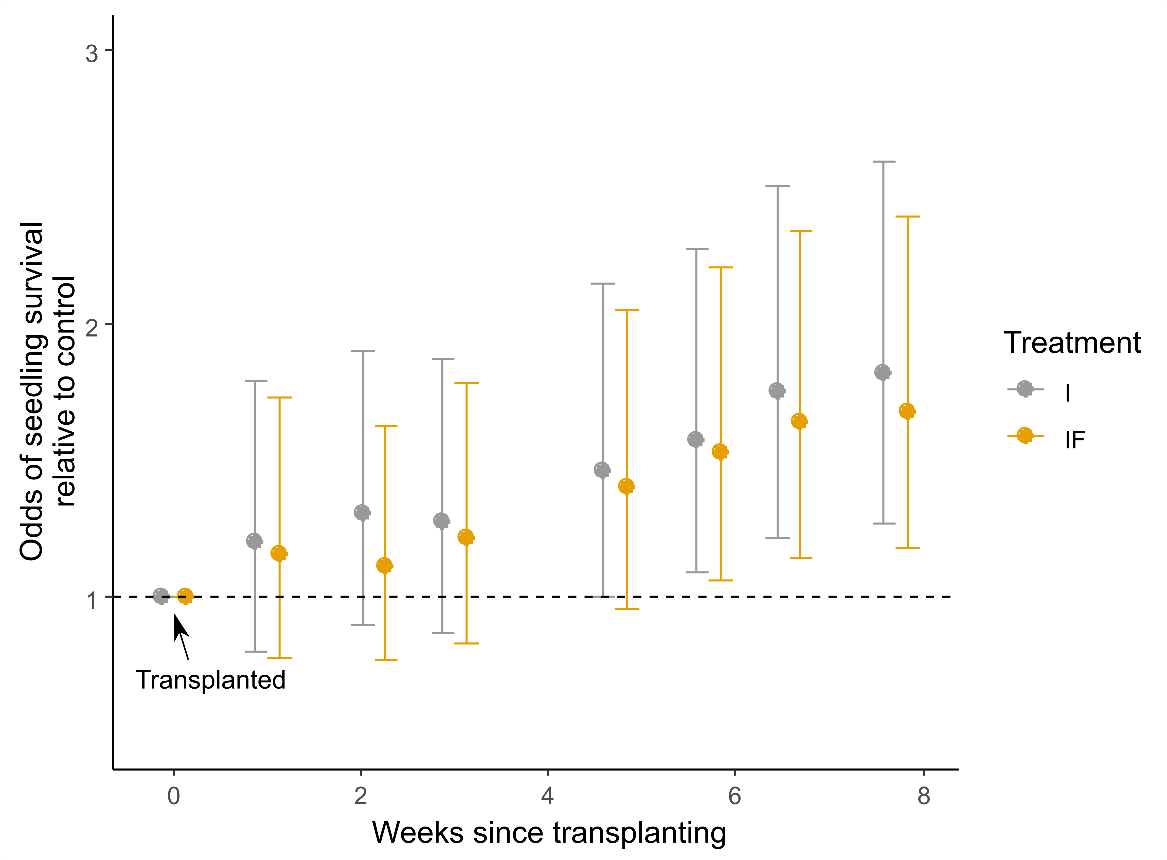

Figure S5: Effects of pesticide treatment (insecticide, I; insecticide + fungicide, IF) on individual seedling survival, relative to control seedlings (± 95% C.I.). The dashed horizontal line represents survival rates equal to the control group, averaged across all other factors.

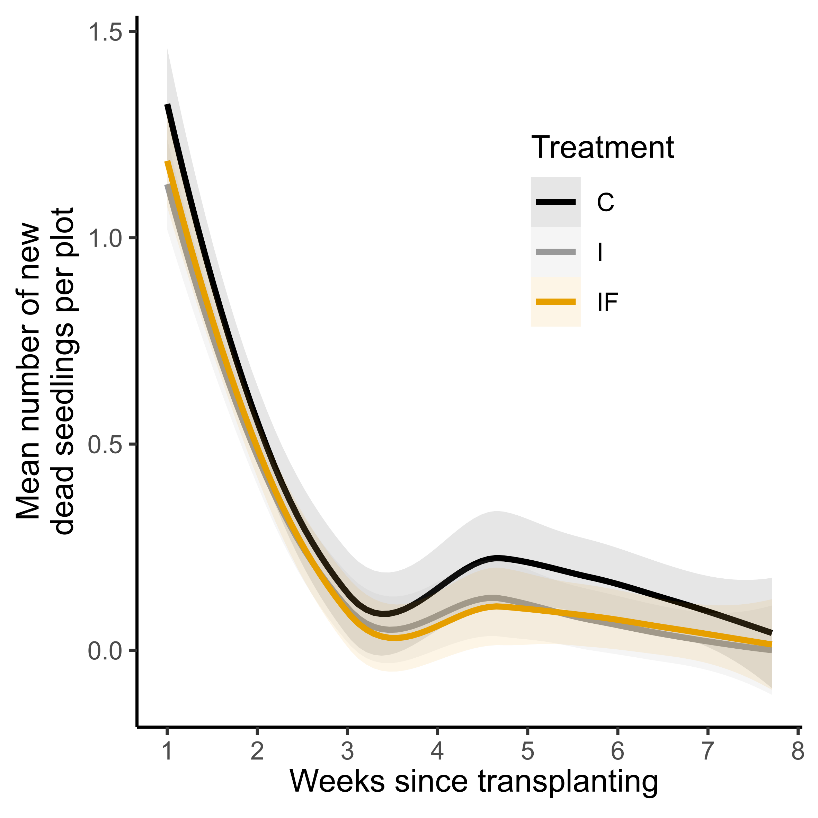

Figure S6: The number of new dead seedlings per plot at each sampling period, split by treatment (control, C; insecticide, I; insecticide + fungicide, IF). Lines are smoothed loess fits ± 95% C.I.

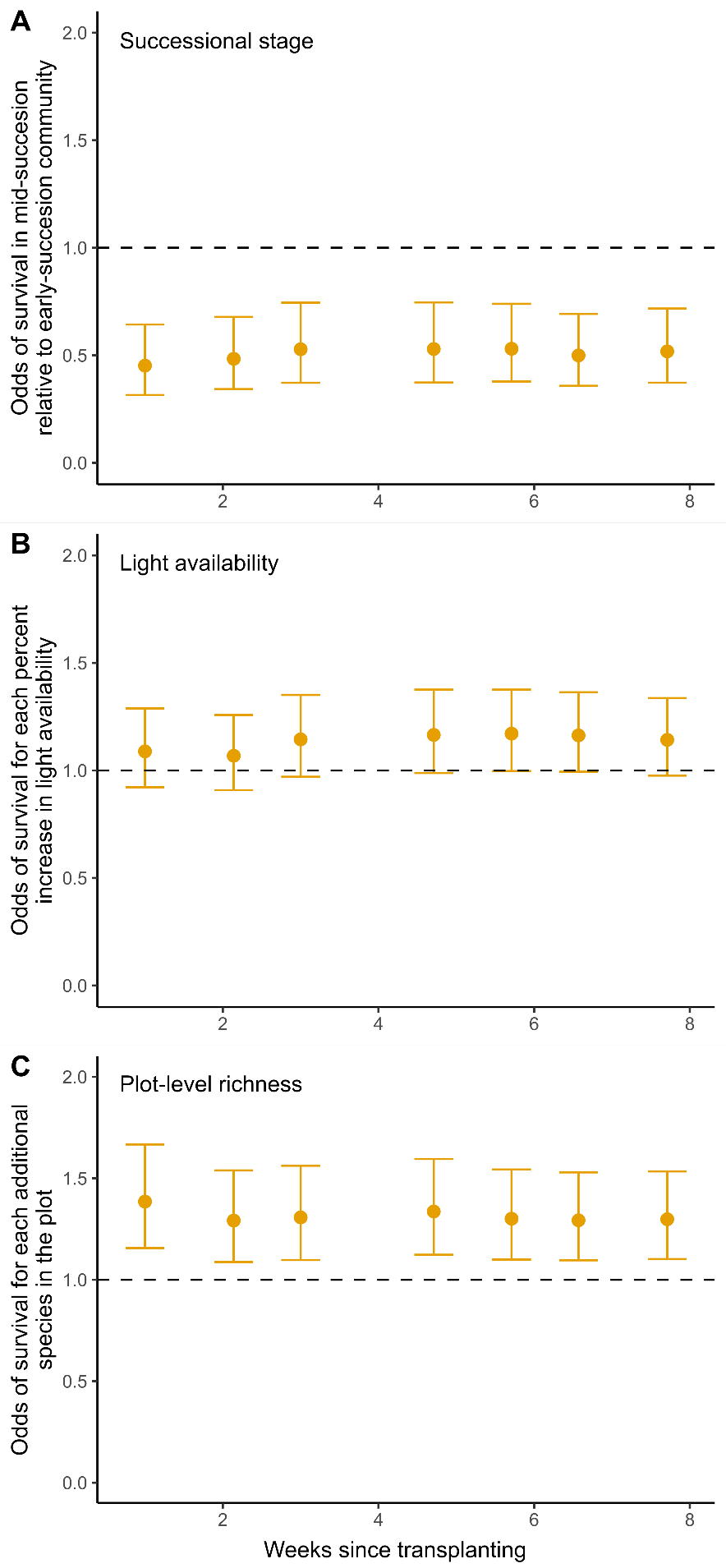

Figure S7: The effect of (a) successional stage, (b) light availability, and (c) species richness, in determining the odds of survival of planted seedlings, accounting for all other factors in the model. Dots and error bars are mean estimates ± 95% C.I.

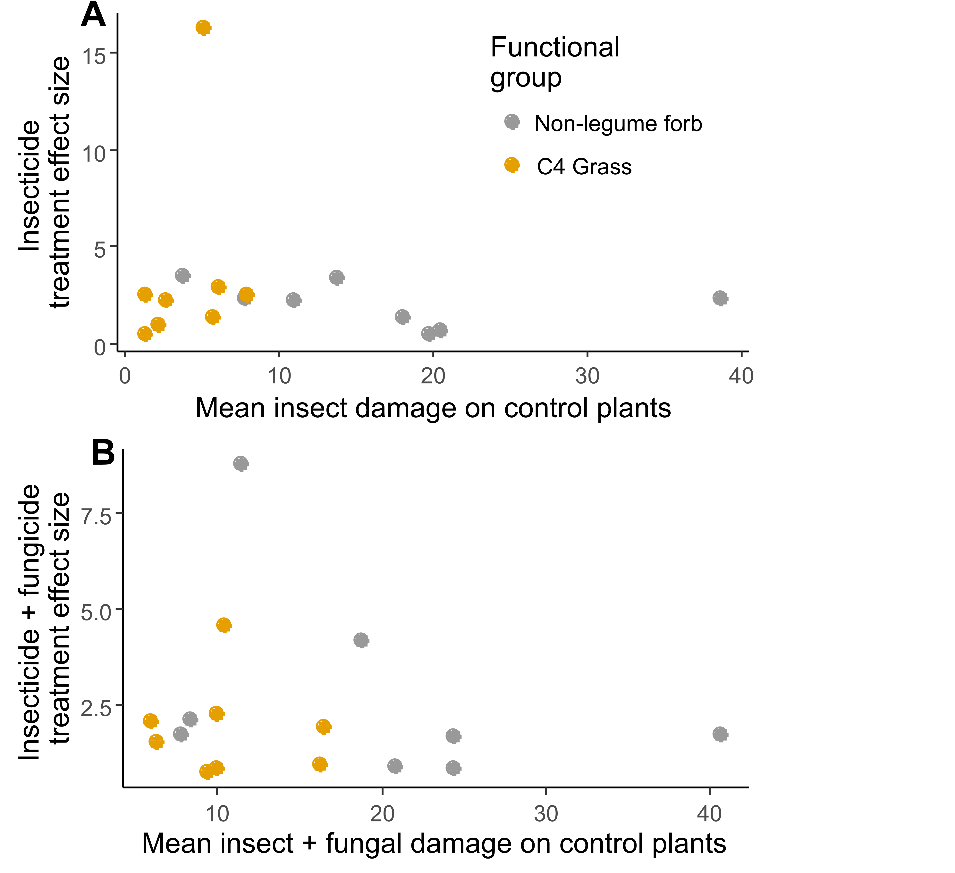

Figure S8: Relationship between effect sizes for (a) Insecticide treatment, (b) Insecticide + Fungicide treatment, and damage rates on control plants.

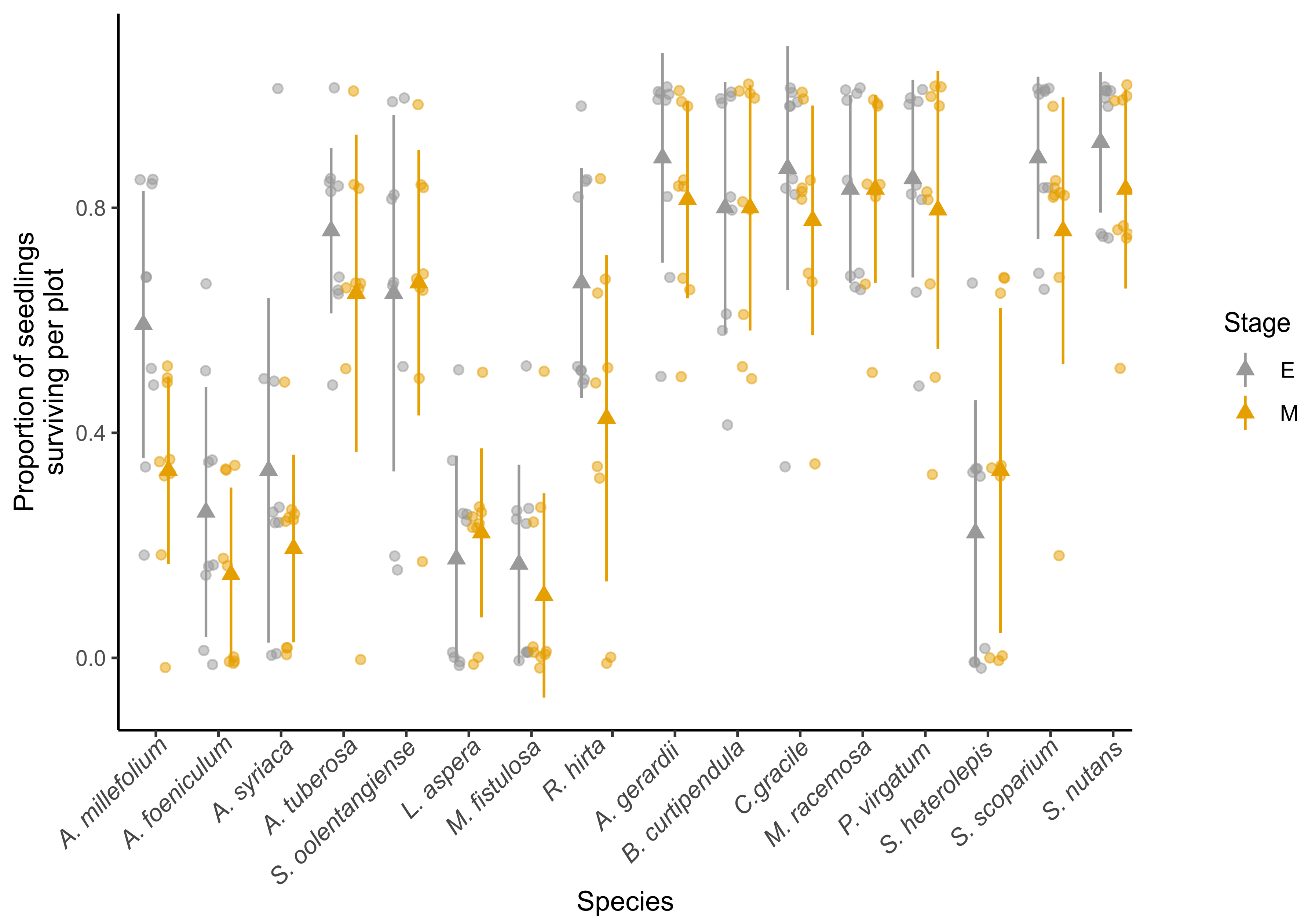

Figure S9: Proportion of seedlings of each species surviving per plot, separated by successional stage (early, E; mid, M) and averaging across all other factors. Dots are individual plots, with triangles and errors bars as mean ± 1 S.D.

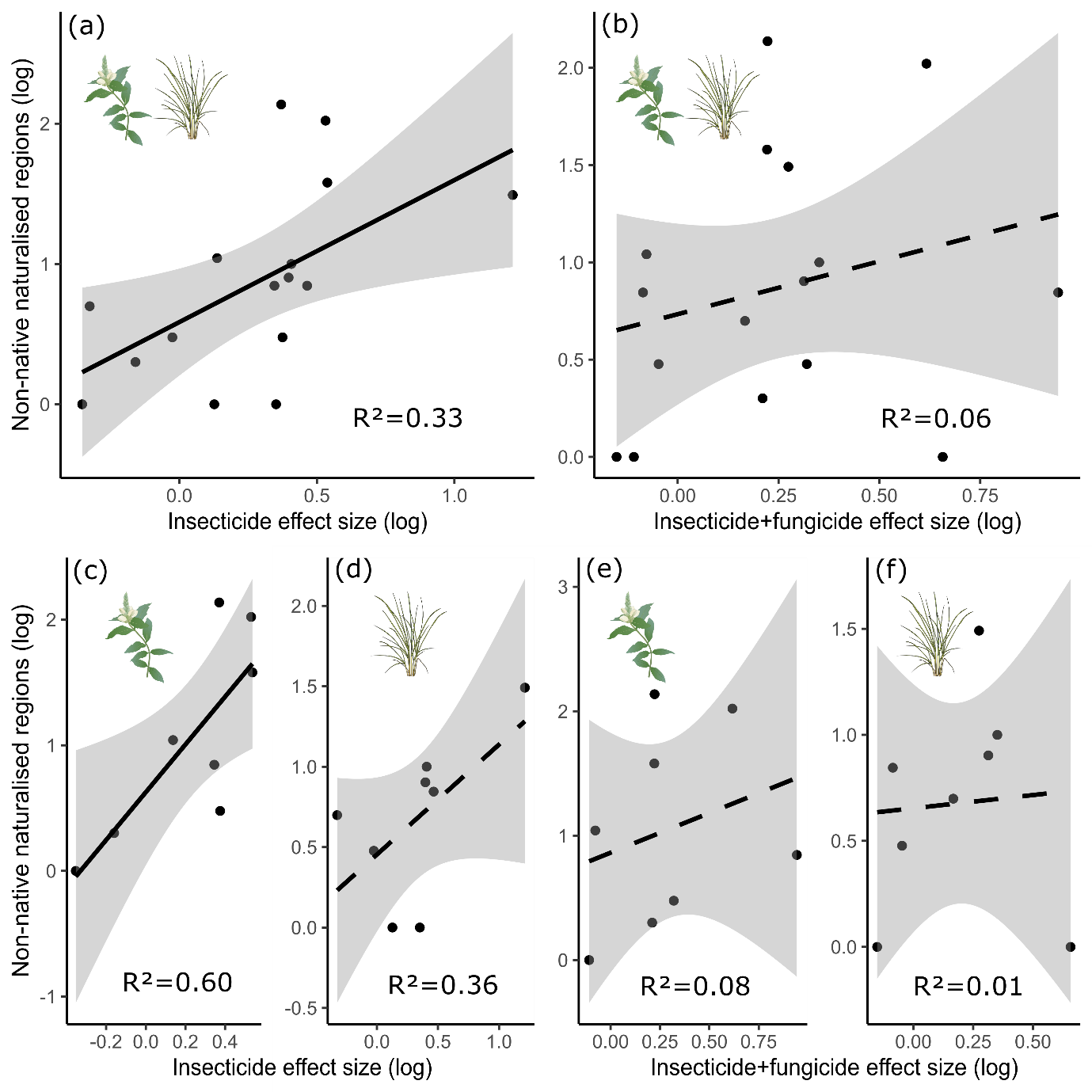

Figure S10: Relationship between treatment effect sizes and the number of regions globally that each species is reported as being naturalised in (from the GloNAF database, van Kleunen et al. 2019), including the outlier *A. gerardii*. Relationships are shown for insecticide effect sizes (a, c, d) and insecticide+fungicide effect sizes (b, e, f), for both forbs and grasses combined (a, b) as well as for forbs separately (c, e) and grasses separately (d, f). Each dot is an individual species. Solid lines indicate statistical significance, dashed lines indicate non-significance.

**Supporting Information: Tables**

*Model Set 1a: Quasibinomial models exploring proportional seedling survival at the plot level*

Table S1: *June 22^nd^*: Log-ratio χ^2^ tests for the main effect of each factor, assuming all other factors are in the model. Interactions were tested and all found to be non-significant. Bold p-values indicate significance at a Bonferroni-corrected α=0.007, stars (*) indicate significance at α=0.05. ‘d.f.’ = degrees of freedom.

| **Factor** | **χ^2^** | **d.f.** | **p-value** |
| --- | --- | --- | --- |
| Treatment | 1.429 | 2 | 0.489 |
| Species | 177.891 | 15 | **<0.001** |
| awMPD | 0.015 | 1 | 0.902 |
| Successional stage | 19.820 | 1 | **<0.001** |
| Light availability | 0.997 | 1 | 0.318 |
| Moisture | 0.701 | 1 | 0.402 |
| Plot richness | 12.608 | 1 | **<0.001** |

R^2^ of model: 0.332

Table S2: *June 30^th^*: Log-ratio χ^2^ tests for the main effect of each factor, assuming all other factors are in the model. Interactions were tested and all found to be non-significant. Bold p-values indicate significance at a Bonferroni-corrected α=0.007, stars (*) indicate significance at α=0.05. ‘d.f.’ = degrees of freedom.

| **Factor** | **χ^2^** | **d.f.** | **p-value** |
| --- | --- | --- | --- |
| Treatment | 2.103 | 2 | 0.349 |
| Species | 193.493 | 15 | **<0.001** |
| awMPD | 0.062 | 1 | 0.803 |
| Successional stage | 17.981 | 1 | **<0.001** |
| Light availability | 0.640 | 1 | 0.424 |
| Moisture | 0.007 | 1 | 0.936 |
| Plot richness | 8.586 | 1 | **0.003** |

R^2^ of model: 0.339

Table S3: *July 6^th^*: Log-ratio χ^2^ tests for the main effect of each factor, assuming all other factors are in the model. Interactions were tested and all found to be non-significant. Bold p-values indicate significance at a Bonferroni-corrected α=0.007, stars (*) indicate significance at α=0.05. ‘d.f.’ = degrees of freedom.

| **Factor** | **χ^2^** | **d.f.** | **p-value** |
| --- | --- | --- | --- |
| Treatment | 2.048 | 2 | 0.359 |
| Species | 190.933 | 15 | **<0.001** |
| awMPD | 0.232 | 1 | 0.630 |
| Successional stage | 13.327 | 1 | **<0.001** |
| Light availability | 2.57 | 1 | 0.109 |
| Moisture | 0.018 | 1 | 0.894 |
| Plot richness | 9.084 | 1 | **0.003** |

R^2^ of model: 0.343

Table S4: *July 18^th^*: Log-ratio χ^2^ tests for the main effect of each factor, assuming all other factors are in the model. Interactions were tested and all found to be non-significant. Bold p-values indicate significance at a Bonferroni-corrected α=0.007, stars (*) indicate significance at α=0.05. ‘d.f.’ = degrees of freedom.

| **Factor** | **χ^2^** | **d.f.** | **p-value** |
| --- | --- | --- | --- |
| Treatment | 5.182 | 2 | 0.075 |
| Species | 200.233 | 15 | **<0.001** |
| awMPD | 0.679 | 1 | 0.410 |
| Successional stage | 13.329 | 1 | **<0.001** |
| Light availability | 3.283 | 1 | 0.070 |
| Moisture | 0.092 | 1 | 0.762 |
| Plot richness | 10.738 | 1 | **0.001** |

R^2^ of model: 0.363

Table S5: *July 25^th^*: Log-ratio χ^2^ tests for the main effect of each factor, assuming all other factors are in the model. Interactions were tested and all found to be non-significant. Bold p-values indicate significance at a Bonferroni-corrected α=0.007, stars (*) indicate significance at α=0.05. ‘d.f.’ = degrees of freedom.

| **Factor** | **χ^2^** | **d.f.** | **p-value** |
| --- | --- | --- | --- |
| Treatment | 8.157 | 2 | 0.017* |
| Species | 225.196 | 15 | **<0.001** |
| awMPD | 0.879 | 1 | 0.348 |
| Successional stage | 14.112 | 1 | **<0.001** |
| Light availability | 3.708 | 1 | 0.054 |
| Moisture | 0.531 | 1 | 0.466 |
| Plot richness | 9.487 | 1 | **0.002** |

R^2^ of model: 0.381

Table S6: *July 31^st^* Log-ratio χ^2^ tests for the main effect of each factor, assuming all other factors are in the model. Interactions were tested and all found to be non-significant. Bold p-values indicate significance at a Bonferroni-corrected α=0.007, stars (*) indicate significance at α=0.05. ‘d.f.’ = degrees of freedom.

| **Factor** | **χ^2^** | **d.f.** | **p-value** |
| --- | --- | --- | --- |
| Treatment | 11.660 | 2 | **0.003** |
| Species | 226.920 | 15 | **<0.001** |
| awMPD | 1.175 | 1 | 0.278 |
| Successional stage | 17.497 | 1 | **<0.001** |
| Light availability | 3.537 | 1 | 0.060 |
| Moisture | 0.127 | 1 | 0.721 |
| Plot richness | 9.371 | 1 | **0.002** |

R^2^ of model: 0.382

Table S7: *August 8^th^*: Log-ratio χ^2^ tests for the main effect of each factor, assuming all other factors are in the model. Interactions were tested and all found to be non-significant. Bold p-values indicate significance at a Bonferroni-corrected α=0.007, stars (*) indicate significance at α=0.05. ‘d.f.’ = degrees of freedom.

| **Factor** | **χ^2^** | **d.f.** | **p-value** |
| --- | --- | --- | --- |
| Treatment | 13.264 | 2 | **0.001** |
| Species | 225.037 | 15 | **<0.001** |
| awMPD | 0.861 | 1 | 0.354 |
| Successional stage | 15.870 | 1 | **<0.001** |
| Light availability | 2.767 | 1 | 0.096 |
| Moisture | 0.190 | 1 | 0.663 |
| Plot richness | 9.787 | 1 | **0.002** |

R^2^ of model: 0.382

*Model Set 1b: Poisson model exploring number of new dead per plot at each time point*

Table S8: Log-ratio χ^2^ tests for the effect of each factor, assuming all other factors are in the model. Bold p-values indicate significance at α=0.05. ‘d.f.’ = degrees of freedom.

| **Factor** | **χ^2^** | **d.f.** | **p-value** |
| --- | --- | --- | --- |
| Treatment | 2.070 | 2 | 0.355 |
| Time | 207.534 | 1 | **<0.001** |
| Species | 71.820 | 15 | **<0.001** |
| awMPD | 0.944 | 1 | 0.331 |
| Successional stage | 9.895 | 1 | **0.002** |
| Light availability | 1.065 | 1 | 0.302 |
| Moisture | 0.011 | 1 | 0.917 |
| Plot richness | 5.270 | 1 | **0.022** |
| Treatment*Time | 14.107 | 2 | **0.001** |

*Model Set 2: Binomial models exploring individual seedling survival*

Table S9: *June 22^nd^*: Log-ratio χ^2^ tests for the main effect of each factor, assuming all other factors are in the model. Interactions were tested and all found to be non-significant. Bold p-values indicate significance at a Bonferroni-corrected α=0.007, stars (*) indicate significance at α=0.05. ‘d.f.’ = degrees of freedom.

| **Factor** | **χ^2^** | **d.f.** | **p-value** |
| --- | --- | --- | --- |
| Treatment | 0.923 | 2 | 0.631 |
| Successional stage | 10.355 | 1 | **0.001** |
| Initial height | 34.384 | 1 | **<0.001** |

Table S10: *June 30^th^*: Log-ratio χ^2^ tests for the main effect of each factor, assuming all other factors are in the model. Interactions were tested and all found to be non-significant. Bold p-values indicate significance at a Bonferroni-corrected α=0.007, stars (*) indicate significance at α=0.05. ‘d.f.’ = degrees of freedom.

| **Factor** | **χ^2^** | **d.f.** | **p-value** |
| --- | --- | --- | --- |
| Treatment | 1.952 | 2 | 0.377 |
| Successional stage | 12.860 | 1 | **<0.001** |
| Initial height | 47.066 | 1 | **<0.001** |

Table S11: *July 6^th^*: Log-ratio χ^2^ tests for the main effect of each factor, assuming all other factors are in the model. Interactions were tested and all found to be non-significant. Bold p-values indicate significance at a Bonferroni-corrected α=0.007, stars (*) indicate significance at α=0.05. ‘d.f.’ = degrees of freedom.

| **Factor** | **χ^2^** | **d.f.** | **p-value** |
| --- | --- | --- | --- |
| Treatment | 1.755 | 2 | 0.416 |
| Successional stage | 8.681 | 1 | **0.003** |
| Initial height | 40.656 | 1 | **<0.001** |

Table S12: *July 18^th^*: Log-ratio χ^2^ tests for the main effect of each factor, assuming all other factors are in the model. Interactions were tested and all found to be non-significant. Bold p-values indicate significance at a Bonferroni-corrected α=0.007, stars (*) indicate significance at α=0.05. ‘d.f.’ = degrees of freedom.

| **Factor** | **χ^2^** | **d.f.** | **p-value** |
| --- | --- | --- | --- |
| Treatment | 4.675 | 2 | 0.097 |
| Successional stage | 7.905 | 1 | **0.005** |
| Initial height | 39.618 | 1 | **<0.001** |

Table S13: *July 25^th^*: Log-ratio χ^2^ tests for the main effect of each factor, assuming all other factors are in the model. Interactions were tested and all found to be non-significant. Bold p-values indicate significance at a Bonferroni-corrected α=0.007, stars (*) indicate significance at α=0.05. ‘d.f.’ = degrees of freedom.

| **Factor** | **χ^2^** | **d.f.** | **p-value** |
| --- | --- | --- | --- |
| Treatment | 7.439 | 2 | 0.024* |
| Successional stage | 8.338 | 1 | **0.004** |
| Initial height | 39.630 | 1 | **<0.001** |

Table S14: *July 31^st^*: Log-ratio χ^2^ tests for the main effect of each factor, assuming all other factors are in the model. Interactions were tested and all found to be non-significant. Bold p-values indicate significance at a Bonferroni-corrected α=0.007, stars (*) indicate significance at α=0.05. ‘d.f.’ = degrees of freedom.

| **Factor** | **χ^2^** | **d.f.** | **p-value** |
| --- | --- | --- | --- |
| Treatment | 11.256 | 2 | **0.004** |
| Successional stage | 11.863 | 1 | **0.001** |
| Initial height | 34.991 | 1 | **<0.001** |

Table S15: *August 8^th^*: Log-ratio χ^2^ tests for the main effect of each factor, assuming all other factors are in the model. Interactions were tested and all found to be non-significant. Bold p-values indicate significance at a Bonferroni-corrected α=0.007, stars (*) indicate significance at α=0.05. ‘d.f.’ = degrees of freedom.

| **Factor** | **χ^2^** | **d.f.** | **p-value** |
| --- | --- | --- | --- |
| Treatment | 12.948 | 2 | **0.002** |
| Successional stage | 9.953 | 1 | **0.002** |
| Initial height | 36.520 | 1 | **<0.001** |

*Model Set 3: Binomial models exploring individual survival per species at time 7 only.*

Note treatment is generally not significant, likely due to high variation and low within-species replication.

Table S16: *Achillea millefolium*: Log-ratio χ^2^ tests for the main effect of each factor, assuming all other factors are in the model. Interactions were tested and all found to be non-significant. Bold p-values indicate significance at α=0.05. ‘d.f.’ = degrees of freedom.

| **Factor** | **χ^2^** | **d.f.** | **p-value** |
| --- | --- | --- | --- |
| Treatment | 2.912 | 2 | 0.672 |
| Successional stage | 7.388 | 1 | **0.007** |
| Initial height | 0.156 | 1 | 0.693 |

Table S17: *Agastache foeniculum*: Log-ratio χ^2^ tests for the main effect of each factor, assuming all other factors are in the model. Interactions were tested and all found to be non-significant. Bold p-values indicate significance at α=0.05. ‘d.f.’ = degrees of freedom.

| **Factor** | **χ^2^** | **d.f.** | **p-value** |
| --- | --- | --- | --- |
| Treatment | 0.558 | 2 | 0.756 |
| Successional stage | 1.824 | 1 | 0.179 |
| Initial height | 0.303 | 1 | 0.582 |

Table S18: *Andropogon gerardii*: Log-ratio χ^2^ tests for the main effect of each factor, assuming all other factors are in the model. Interactions were tested and all found to be non-significant. Bold p-values indicate significance at α=0.05. ‘d.f.’ = degrees of freedom.

| **Factor** | **χ^2^** | **d.f.** | **p-value** |
| --- | --- | --- | --- |
| Treatment | 2.279 | 2 | 0.320 |
| Successional stage | 0.907 | 1 | 0.340 |
| Initial height | 4.943 | 1 | **0.026** |

Table S19: *Asclepias syriaca*: Log-ratio χ^2^ tests for the main effect of each factor, assuming all other factors are in the model. Interactions were tested and all found to be non-significant. Bold p-values indicate significance at α=0.05. ‘d.f.’ = degrees of freedom.

| **Factor** | **χ^2^** | **d.f.** | **p-value** |
| --- | --- | --- | --- |
| Treatment | 2.996 | 2 | 0.224 |
| Successional stage | 1.494 | 1 | 0.222 |
| Initial height | 0.458 | 1 | 0.499 |

Table S20: *Asclepias tuberosa*: Log-ratio χ^2^ tests for the main effect of each factor, assuming all other factors are in the model. Interactions were tested and all found to be non-significant. Bold p-values indicate significance at α=0.05. ‘d.f.’ = degrees of freedom.

| **Factor** | **χ^2^** | **d.f.** | **p-value** |
| --- | --- | --- | --- |
| Treatment | 3.021 | 2 | 0.221 |
| Successional stage | 0.878 | 1 | 0.349 |
| Initial height | 3.744 | 1 | 0.053 |

Table S21: S*ymphyotrichum oolentangiense*: Log-ratio χ^2^ tests for the main effect of each factor, assuming all other factors are in the model. Interactions were tested and all found to be non-significant. Bold p-values indicate significance at α=0.05. ‘d.f.’ = degrees of freedom.

| **Factor** | **χ^2^** | **d.f.** | **p-value** |
| --- | --- | --- | --- |
| Treatment | 1.099 | 2 | 0.577 |
| Successional stage | 0.013 | 1 | 0.911 |
| Initial height | 2.075 | 1 | 0.150 |

Table S22: *Bouteloua curtipendula*: Log-ratio χ^2^ tests for the main effect of each factor, assuming all other factors are in the model. Interactions were tested and all found to be non-significant. Bold p-values indicate significance at α=0.05. ‘d.f.’ = degrees of freedom.

| **Factor** | **χ^2^** | **d.f.** | **p-value** |
| --- | --- | --- | --- |
| Treatment | 1.906 | 2 | 0.386 |
| Successional stage | 0.549 | 1 | 0.459 |
| Initial height | 13.281 | 1 | **<0.001** |

Table S23: *Chondrosum gracile*: Log-ratio χ^2^ tests for the main effect of each factor, assuming all other factors are in the model. Interactions were tested and all found to be non-significant. Bold p-values indicate significance at α=0.05. ‘d.f.’ = degrees of freedom.

| **Factor** | **χ^2^** | **d.f.** | **p-value** |
| --- | --- | --- | --- |
| Treatment | 1.672 | 2 | 0.433 |
| Successional stage | 1.032 | 1 | 0.310 |
| Initial height | 3.492 | 1 | 0.062 |

Table S24: *Liatris aspera*: Log-ratio χ^2^ tests for the main effect of each factor, assuming all other factors are in the model. Interactions were tested and all found to be non-significant. Bold p-values indicate significance at α=0.05. ‘d.f.’ = degrees of freedom.

| **Factor** | **χ^2^** | **d.f.** | **p-value** |
| --- | --- | --- | --- |
| Treatment | 1.167 | 2 | 0.558 |
| Successional stage | 0.047 | 1 | 0.828 |
| Initial height | 6.713 | 1 | **0.010** |

Table S25: *Monarda fistulosa*: Log-ratio χ^2^ tests for the main effect of each factor, assuming all other factors are in the model. Interactions were tested and all found to be non-significant. Bold p-values indicate significance at α=0.05. ‘d.f.’ = degrees of freedom.

| **Factor** | **χ^2^** | **d.f.** | **p-value** |
| --- | --- | --- | --- |
| Treatment | 4.089 | 2 | 0.129 |
| Successional stage | 0.903 | 1 | 0.342 |
| Initial height | 8.391 | 1 | **0.004** |

Table S26: *Muhlenbergia racemosa*: Log-ratio χ^2^ tests for the main effect of each factor, assuming all other factors are in the model. Interactions were tested and all found to be non-significant. Bold p-values indicate significance at α=0.05. ‘d.f.’ = degrees of freedom.

| **Factor** | **χ^2^** | **d.f.** | **p-value** |
| --- | --- | --- | --- |
| Treatment | 2.413 | 2 | 0.299 |
| Successional stage | 0.043 | 1 | 0.837 |
| Initial height | 0.468 | 1 | 0.439 |

Table S27: *Panicum virgatum*: Log-ratio χ^2^ tests for the main effect of each factor, assuming all other factors are in the model. Interactions were tested and all found to be non-significant. Bold p-values indicate significance at α=0.05. ‘d.f.’ = degrees of freedom.

| **Factor** | **χ^2^** | **d.f.** | **p-value** |
| --- | --- | --- | --- |
| Treatment | 5.775 | 2 | 0.056 |
| Successional stage | 0.502 | 1 | 0.478 |
| Initial height | 0.196 | 1 | 0.658 |

Table S28: *Rudbeckia hirta*: Log-ratio χ^2^ tests for the main effect of each factor, assuming all other factors are in the model. Interactions were tested and all found to be non-significant. Bold p-values indicate significance at α=0.05. ‘d.f.’ = degrees of freedom.

| **Factor** | **χ^2^** | **d.f.** | **p-value** |
| --- | --- | --- | --- |
| Treatment | 8.387 | 2 | **0.015** |
| Successional stage | 6.395 | 1 | **0.011** |
| Initial height | 2.804 | 1 | 0.094 |

Table S29: *Schizachyrium scoparium*: Log-ratio χ^2^ tests for the main effect of each factor, assuming all other factors are in the model. Interactions were tested and all found to be non-significant. Bold p-values indicate significance at α=0.05. ‘d.f.’ = degrees of freedom.

| **Factor** | **χ^2^** | **d.f.** | **p-value** |
| --- | --- | --- | --- |
| Treatment | 2.099 | 2 | 0.350 |
| Successional stage | 2.502 | 1 | 0.113 |
| Initial height | 0.002 | 1 | 0.967 |

Table S30: *Sorghastrum nutans*: Log-ratio χ^2^ tests for the main effect of each factor, assuming all other factors are in the model. Interactions were tested and all found to be non-significant. Bold p-values indicate significance at α=0.05. ‘d.f.’ = degrees of freedom.

| **Factor** | **χ^2^** | **d.f.** | **p-value** |
| --- | --- | --- | --- |
| Treatment | 0.014 | 2 | 0.993 |
| Successional stage | 0.787 | 1 | 0.375 |
| Initial height | 0.176 | 1 | 0.675 |

Table S31: *Sporobolus heterolepis*: Log-ratio χ^2^ tests for the main effect of each factor, assuming all other factors are in the model. Interactions were tested and all found to be non-significant. Bold p-values indicate significance at α=0.05. ‘d.f.’ = degrees of freedom.

| **Factor** | **χ^2^** | **d.f.** | **p-value** |
| --- | --- | --- | --- |
| Treatment | 0.674 | 2 | 0.714 |
| Successional stage | 0.705 | 1 | 0.401 |
| Initial height | 0.269 | 1 | 0.604 |

Table S32: Relationship between damage on control plants and effect sizes.

| **Test** | **t-value** | **Degrees of freedom** | **p-value** |
| --- | --- | --- | --- |
| Effect size I ~ Insect damage on control plants | -0.614 | 15 | 0.549 |
| Effect size IF ~ Fungal damage on control plants | -0.135 | 15 | 0.374 |
| Effect size IF ~ Insect + fungal damage on control plants | -0.564 | 15 | 0.582 |

Table S33: Relationship between plant traits and effect sizes. Effect sizes were logged for analysis, and functional group was included as a random variable. Bold p-values indicate significance at α=0.05.

| **Test** | **β coefficient** | **χ^2^** | **d.f.** | **p-value** |
| --- | --- | --- | --- | --- |
| *Aboveground Principal Component traits* | | | | |
| Effect size I ~ PC1 | -0.127 | 4.049 | 1 | **0.044** |
| Effect size I ~ PC2 | -0.119 | 1.398 | 1 | 0.237 |
| Effect size IF ~ PC1 | -0.053 | 1.110 | 1 | 0.292 |
| Effect size IF ~ PC2 | -0.061 | 0.526 | 1 | 0.468 |
| *Individual traits* | | | | |
| Effect size I ~ SLA | 0.021 | 2.902 | 1 | 0.088 |
| Effect size I ~ Leaf N | 0.248 | 2.405 | 1 | 0.121 |
| Effect size IF ~ SLA | 0.013 | 1.499 | 1 | 0.221 |
| Effect size IF ~ Leaf N | 0.128 | 0.927 | 1 | 0.336 |

Table S34: Relationship between effect sizes and number of non-naturalised regions that experimental species are reported in. Bold p-values indicate significance at α=0.05.

| **Test** | **t-value** | **Degrees of freedom** | **p-value** |
| --- | --- | --- | --- |
| *With outlier excluded* | | | |
| Naturalised regions ~ Overall effect size I | 2.935 | 13 | **0.012** |
| Naturalised regions ~ Overall effect size IF | 1.692 | 13 | 0.115 |
| Naturalised regions ~ Effect size I (forbs only) | 3.015 | 6 | **0.024** |
| Naturalised regions ~ Effect size IF (forbs only) | 0.709 | 6 | 0.505 |
| Naturalised regions ~ Effect size I (grasses only) | 2.396 | 5 | 0.062 |
| Naturalised regions ~ Effect size IF (grasses only) | 2.486 | 5 | 0.055 |
| *With outlier included* | | | |
| Naturalised regions ~ Overall effect size I | 2.622 | 14 | **0.020** |
| Naturalised regions ~ Overall effect size IF | 0.955 | 14 | 0.355 |
| Naturalised regions ~ Effect size I (forbs only) | 3.015 | 6 | **0.024** |
| Naturalised regions ~ Effect size IF (forbs only) | 0.709 | 6 | 0.505 |
| Naturalised regions ~ Effect size I (grasses only) | 1.853 | 6 | 0.113 |
| Naturalised regions ~ Effect size IF (grasses only) | 0.163 | 6 | 0.876 |
